# Humans homozygous for rare or common hypomorphic *IL23R* variants are prone to tuberculosis

**DOI:** 10.64898/2026.03.23.713554

**Authors:** Diana Olguín Calderón, Laura E. Kilpatrick, Clément Conil, Quentin Philippot, Masato Ogishi, Joseph Vellutini, Ji Eun Han, Narelle Keating, Hailun Li, Geetha Rao, Jonathan Bohlen, Charles S. Lay, Simon Platt, Gaspard Kerner, Elsa Feredj, Jessica N. Peel, Mana Momenilandi, Yoann Seeleuthner, Candice Lainé, Camille Soudée, Claire Leloup, Cecile Debuisson, Fanny Lanternier, Samuel Bitoun, Stephan Pavy, Xavier Mariette, Aniss Rafik, Hanaa Skhoun, Hanane El Ouazzani, Ismael Abderahmani-Ghorfi, Jamila El-Baghdadi, Andrés Baena, Manuela Tejada-Giraldo, Luis FernandoBarrera, Andrés Augusto Arias, Giovanna Fabio, Maria Carrabba, Melike Emiroglu, Liliana Bezrodnik, Loubna El Zein, Hassan Hammoud, Peter K. Gregersen, Benjamin Terrier, Rafael Leon Lopez, Marion Touzet, Vincent Pestre, Marlène Pasquet, Lars Rogge, Michael Fayon, François Galode, Eric Jeziorski, Darragh Duffy, Lluis Quintana-Murci, Etienne Patin, Charlotte Cunningham-Rundles, Isabelle Meyts, Shen-Ying Zhang, Qian Zhang, Emmanuelle Jouanguy, Bertrand Boisson, Jérémie Rosain, Vivien Béziat, Mohammad Shahrooei, Seyed Alireza Mahdaviani, Nima Rezaei, Nima Parvaneh, Zahra Chavoshzadeh, Niloufar Yazdanpanah, Nathalie Aladjidi, Antoni Noguera-Julian, Ana Esteve-Solé, Laia-Alsina, Davood Mansouri, Sevgi Keles, Mediha Gonenc Ortakoylu, Deniz Aygun, Esra Yucel, Ayca Kiykim, Yildiz Camcioglu, Cindy S. Ma, Stuart G. Tangye, Peng Zhang, Laurent Abel, Peter D. Craggs, Jean-Laurent Casanova, Aurélie Cobat, Anne Puel, Jacinta Bustamante, Stephen J. Hill, Stéphanie Boisson-Dupuis

**Affiliations:** Laboratory of Human Genetics of Infectious Diseases, Necker Branch, INSERM U1163, Necker Hospital for Sick Children, Paris, France; Imagine Institute, Paris Cité University, Paris, France; Division of Biomolecular Science and Medicinal Chemistry, School of Pharmacy, University of Nottingham, Nottingham, UK; Centre of Membrane Proteins and Receptors, University of Birmingham and University of Nottingham, The Midlands, Nottingham, UK; St Giles Laboratory of Human Genetics of Infectious Diseases, Rockefeller Branch, Rockefeller University, New York, NY, USA; Garvan Institute of Medical Research, Darlinghurst, Australia; Faculty of Medicine and Health, School of Clinical Medicine, University of New South Wales Sydney, Kensington, Australia; Gene Center and Department of Biochemistry, Ludwig-Maximilians-Universität, Munich, Germany; Department of Pediatrics, Dr. von Hauner Childrens Hospital, LMU Klinikum, Munich, Germany; German Center for Child and Adolescent Health (DZKJ), Munich, Germany; Division of Physiology, Pharmacology and Neuroscience, School of Life Sciences, University of Nottingham, Nottingham, UK; Human Evolutionary Genetics Unit, Institut Pasteur, Paris Cité University, CNRS UMR2000, Paris, France; Immunoregulation Unit, Department of Immunology, Institut Pasteur, Paris Cité University, Paris, France; General Pediatrics Department, Hôpital des Enfants, Centre Hospitalier Universitaire de Toulouse, Toulouse, France; Service de Maladies Infectieuses et Tropicales, Hôpital Necker-Enfants Malades, Assistance Publique-Hôpitaux de Paris (AP-HP), Paris, France; Translational Mycology Research Group, Mycology Department, National Reference Center for Invasive Mycoses and Antifungals, Institut Pasteur, Paris Cité University, Paris, France; Department of Rheumatology, AP-HP, Hôpital Bicêtre, Université Paris-Saclay, INSERM, CEA, UMR 1184, FHU CARE, Le Kremlin Bicêtre, France; Genetics Unit, Mohamed V Military Hospital, Hay Riad, Morocco; Department of Pulmonology, Military Hospital Mohammed V, Medical and Pharmacy School of Rabat, Mohammed V University, Rabat, Morocco; Facultad de Medicina, Grupo de Inmunología Celular e Inmunogenética (GICIG), Universidad de Antioquia (UdeA), Medellín, Colombia; Departamento de Microbiología y Parasitología, Facultad de Medicina, UdeA, Medellín, Colombia; Inborn Errors of Immunity Group, Department of Microbiology and Parasitology, School of Medicine, UdeA, Medellín, Colombia; Instituto de Investigaciones Médicas, UdeA, Medellín, Colombia; School of Microbiology, UdeA, Medellín, Colombia; Department of Internal Medicine, Fondazione Istituto di Ricovero e Cura a Carattere Scientifico (IRCCS), Ca’ Granda Ospedale Maggiore Policlinico, Milan, Italy; Division of Pediatric Infectious Diseases, Department of Pediatrics, Selcuk University Faculty of Medicine, Konya, Turkey; Grupo de Inmunología-Instituto Multidisciplinario de Investigaciones en Patologías Pediátricas (IMIPP-CONICET), Hospital de Niños “Dr. Ricardo Gutierrez”, Buenos Aires, Argentina; Center for Clinical Immunology, Buenos Aires, Argentina; Biology Department, Lebanese University, Beirut, Lebanon; Saint George Hospital, Beirut, Lebanon; Robert S. Boas Center for Genomics and Human Genetics, Institute of Molecular Medicine, Feinstein Institutes for Medical Research, Manhasset, NY, USA; Department of Internal Medicine, Cochin Hospital, University of Paris, AP-HP, Paris, France; Unidad de Gestión Clínica de Cuidados Intensivos, Instituto Maimónides de Investigación Biomédica de Córdoba (IMIBIC), Hospital Universitario Reina Sofía, Universidad de Córdoba (UCO), Córdoba, Spain; Department of Internal Medicine, Hospital Centre Avignon, Avignon, France; Department of Pediatric Hematology and Oncology, Centre Hospitalo-Universitaire de Toulouse, Toulouse, France; CHU Bordeaux, Département de Pédiatrie, CIC-P INSERM 1401, Bordeaux, France; Centre de Recherche Cardio-thoracique de Bordeaux, Université de Bordeaux, INSERM U1045, Bordeaux, France; Pediatric Pulmonology, Pellegrin Hospital, Bordeaux University Hospital, Bordeaux, France; Centre Constitutif des Maladies Respiratoires Rares de L’enfant (RESPIRARE), Bordeaux, France; Service Urgences Post-urgences Pédiatriques, PCCEI, CeRéMAIA, Univ Montpellier, CHU Montpellier, Montpellier, France; Translational Immunology Unit, Institut Pasteur, Paris Cité University, Paris, France; Single Cell Biomarkers UTechS, Institut Pasteur, Paris Cité University, Paris, France; Department of Medicine, Icahn School of Medicine at Mount Sinai, New York, NY, USA; Department of Pediatrics, Icahn School of Medicine at Mount Sinai, New York, NY, USA; Department of Pediatrics, University Hospitals Leuven, Leuven, Belgium; Department of Microbiology, Immunology and Transplantation, KU Leuven, Leuven, Belgium; Study Center for Primary Immunodeficiencies, Necker Hospital for Sick Children, AP-HP, Paris, France; Pediatric Infections Research Center, Mofid Children’s Hospital, Shahid Beheshti University of Medical Sciences, Tehran, Iran; Pediatric Respiratory Diseases Research Center, National Research Institute of Tuberculosis and Lung Diseases (NRITLD), Shahid Beheshti University of Medical Sciences, Tehran, Iran; Research Center for Immunodeficiencies, Children’s Medical Center, Tehran University of Medical Sciences, Tehran, Iran; Network of Immunity in Infection, Malignancy and Autoimmunity (NIIMA), Universal Scientific Education and Research Network (USERN), Tehran, Iran; Division of Allergy and Clinical Immunology, Department of Pediatrics, Tehran University of Medical Sciences, Tehran, Iran; Pediatric Hemato-Immunology, Pellegrin Hospital, Bordeaux University Hospital, CIC1401, INSERM CICP, Bordeaux, France; Centre de Compétence des Déficits Immunitaires Héréditaires (CEREDIH), Bordeaux, France; Pediatric Infectious Diseases Department, Infectious Diseases and Systemic Inflammatory Response in Pediatrics, Institut de Recerca Sant Joan de Déu (IRSJD), Hospital Sant Joan de Déu, Barcelona, Spain; Centre for Biomedical Network Research on Epidemiology and Public Health (CIBERESP), Madrid, Spain; Departament de Cirurgia i Especialitats Medicoquirúrgiques, Facultat de Medicina I Ciències de La Salut, Universitat de Barcelona, Barcelona, Spain; Study Group for Immune Dysfunction Diseases in Children (GEMDIP), Institut de Recerca Sant Joan de Déu (IRSJD), Barcelona, Spain; Clinical Immunology and Primary Immunodeficiencies Unit, Pediatric Allergy and Clinical Immunology, Department Hospital Sant Joan de Déu, Barcelona, Spain; Department of Clinical Immunology and Infectious Diseases, NRITLD, Shahid Beheshti University of Medical Sciences, Tehran, Iran; The Clinical Tuberculosis and Epidemiology Research Center, NRITLD, Masih Daneshvari Hospital, Shahid Beheshti University of Medical Sciences, Tehran, <u>Iran</u>; Division of Pediatric Allergy and Immunology, Meram Faculty of Medicine, Necmettin Erbakan University, <u>Konya</u>, Turkey; Department of Pulmonary Medicine, Yedikule Chest Diseases and Thoracic Surgery Training and Research Hospital, University of Health Sciences Turkey, Istanbul, Turkey; Department of Paediatric Infectious Diseases, Cerrahpasa Faculty of Medicine, Istanbul Cerrahpasa-University, Istanbul, Turkey; Department of Pediatric Allergy and Immunology, Cerrahpaşa Faculty of Medicine, Istanbul University-Cerrahpaşa, Istanbul, Turkey; Division of Infectious Diseases and Clinical Immunology, Cerrahpasa Faculty of Medicine, Istanbul University-Cerrahpasa, Istanbul, Turkey; Chemical Biology, Medicine Design, <u>GlaxoSmithKline</u>, <u>Stevenage</u>, UK; Crick-GSK Biomedical Linklabs, <u>Medicine Design</u>, <u>GlaxoSmithKline</u>, <u>Stevenage</u>, UK; Department of Pediatrics, Necker Hospital for Sick Children, AP<u>-HP</u>, Paris, France; Howard Hughes Medical Institute, New York, NY, USA

**Author notes:** Correspondence to Stéphanie Boisson-Dupuis, Jean-Laurent Casanova. D. Olguín Calderón, L.E. Kilpatrick, C. Conil, and Q. Philippot contributed equally to this paper. M. Ogishi and J. Vellutini contributed equally to this paper. P.D. Craggs, J.-L. Casanova, A. Cobat, A. Puel, J. Bustamante, S.J. Hill, and S. Boisson-Dupuis contributed equally to this paper.

## Abstract

Homozygosity for rare loss-of-function *IL23R* variants abolishes IL-23–dependent IFN-γ production by lymphocytes, including NK and innate-like T cells, thereby underlying clinical disease due to weakly virulent mycobacterial species. We report selective enrichment in homozygosity for four hypomorphic *IL23R* variants in our cohort of patients with tuberculosis. Three of these *IL23R* alleles are rare (G300V, G149R, and L372F), with a minor allele frequency (MAF) under 1%, but the fourth (R381Q) is surprisingly common, with an MAF as high as 10.2% in certain populations. The other 15 missense alleles found in the homozygous state in public databases are isomorphic. The four hypomorphic IL-23R variants identified dimerize with IL-12Rβ1 and bind IL-23. However, their function is impaired by low levels of cell surface expression (R381Q, G300V) and/or as a consequence of conformational changes altering agonist efficacy. IFN-γ production in response to IL-23 is impaired in innate-like T cells and NK cells. These data suggest that recessive partial IL-23R deficiency, whether due to rare or common variants, confers a predisposition to tuberculosis while preserving immunity to less virulent mycobacteria.

**One sentence summary:** Homozygous hypomorphic *IL23R* variants impair IL-23-dependent IFN-γ production and underlie tuberculosis.

## Introduction

Mendelian susceptibility to mycobacterial diseases (MSMD) is characterized by selective susceptibility to clinical disease due to weakly virulent mycobacteria, such as Bacille Calmette Guérin (BCG) vaccine substrains and environmental mycobacteria (EM) (Boisson-Dupuis et al., 2024; Bustamante, 2020; Casanova and Abel, 2002; Rosain et al., 2018). The genetic etiology of MSMD is known in about half the patients (Casanova and Abel, 2022). Since 1996, 47 genetic disorders involving 22 loci have been reported (Casanova and Abel, 2002, 2022, Rosain et al., 2018, 2023; Bustamante, 2020; Yang et al., 2020; Martin-Fernandez et al., 2022; Bohlen et al., 2023; Philippot et al., 2023; Boisson-Dupuis et al., 2024; Neehus et al., 2024; Casanova, 2025, 2026). All but two of these disorders affect genes involved in IFN-gamma (IFN-γ) production by lymphocytes, responses of mononuclear phagocytes to IFN-γ, or both. CCR2 deficiency affects monocyte migration (Neehus et al., 2024), whereas the mechanism by which inherited ZNFX1 deficiency causes MSMD remains unclear (Le Voyer et al., 2021). The two most common etiologies of MSMD are autosomal recessive (AR) complete IL-12Rβ1 and IL-12p40 deficiencies (Alodayani et al., 2018; Altare et al., 1998a; Altare et al., 1998b; de Beaucoudrey et al., 2010; de Jong et al., 1998; Fieschi et al., 2003; Khavandegar et al., 2024; Melo et al., 2024; Prando et al., 2013). IL-12 is a heterodimer composed of the p35 and p40 subunits that binds to a heterodimeric receptor composed of IL-12Rβ1 and IL-12Rβ2. Remarkably, p40 can also dimerize with p19, forming IL-23, which binds to a heterodimeric receptor composed of IL-12Rβ1 and IL-23R (Oppmann et al., 2000; Parham et al., 2002). Studies of mouse and human IL-12 and IL-23 have suggested that IL-12 is the signature IFN-γ–inducing T_H_1 cytokine, whereas IL-23 is the signature IL-17–inducing T_H_17 cytokine (Acosta-Rodriguez et al., 2007; Cua and Tato, 2010; Fieschi and Casanova, 2003; Teng et al., 2015; Wilson et al., 2007).

However, patients with AR complete IL-23R deficiency present with MSMD (Martinez-Barricarte et al., 2018; Philippot et al., 2023; Staels et al., 2022); the mechanism involves impaired IL-23–dependent IFN-γ production, particularly by MAIT and Vδ2^+^ γδ T cells (Philippot et al., 2023). Chronic mucocutaneous candidiasis (CMC) in patients with complete IL-23R deficiency has incomplete penetrance due to the low contribution of IL-23–induced IL-17A/F-dependent immunity to CMC (Martinez-Barricarte et al., 2018; Philippot et al., 2023; Staels et al., 2022). In parallel, we reported an enrichment in homozygosity for the common *TYK2* P1104A allele in cohorts of patients of European descent with tuberculosis (TB) relative to ethnicity-adjusted controls, and that homozygosity for the *TYK2* P1104A variant selectively impairs, but does not abrogate, the IL-23–dependent induction of IFN-γ (Boisson-Dupuis et al., 2018; Kerner et al., 2019; Kerner et al., 2021). Consistently, we found that impaired IL-23–dependent induction of IFN-γ is the only mechanism of mycobacterial disease common to patients with any of the five forms of AR TYK2 deficiency (Ogishi et al., 2022). We also found that the impaired IL-23–dependent induction of IFN-γ underlies mycobacterial disease in X-linked recessive MCTS1 deficiency (Bohlen et al., 2023) and AR ITK deficiency (Ogishi et al., 2023). Overall, these findings challenge the classical IL-12–T_H_1/IL-23–T_H_17 dichotomy, suggesting that human IL-23 plays an essential role in IFN-γ–mediated host defense against mycobacteria, with complete and partial deficiencies underlying MSMD and TB, respectively. In this context, we hypothesized that other biallelic *IL23R* variants—null or hypomorphic, rare or common—may underlie MSMD or TB.

## Results

### Four hypomorphic biallelic *IL23R* variants

We screened our cohort of 24,046 patients with various infectious diseases, including 1,874 patients with TB (pulmonary and extrapulmonary) and 901 patients with MSMD, and cohorts of individuals from the general population (gnomAD v4.1.0, Analysis Tool for Annotated Variants Data Base (ATAVDB), Great Middle East, Iranome, Turkish Variome, and Browse All Variants Online (BRAVO) variant server) for biallelic coding non-synonymous or essential splice site *IL23R* variants. We identified 19 rare (minor allelic frequency [MAF] < 0.01) and four common (MAF > 0.01) variants (Table 1). Four of these variants (C115Y, c.367+1G>A, E269*, and c.1149-1G>A) had previously been reported to be loss-of-function (LOF) (Martinez-Barricarte et al., 2018; Philippot et al., 2023). We assessed the functional impact of each of the 19 remaining variants—including G300V and V362I, as well as an allele carrying both G300V and V362I—in an overexpression assay. Indeed, individuals homozygous for *IL23R* G300V in our in-house cohort were also homozygous for the *IL23R* V362I variant. We transiently transfected human embryonic kidney 293T (HEK) cells with a plasmid containing the WT *IL12RB1* cDNA under the control of the PGK promoter, together with a plasmid containing either WT or mutant *IL23R* cDNA under the control of a UbC promoter. In contrast to previous reports of *IL23R* LOF variants expressed under the control of strong CMV or EF1a promoters (Martinez-Barricarte et al., 2018; Philippot et al., 2023; Staels et al., 2022), we used the weaker UbC promoter to facilitate assessment of the impact of hypomorphic variants (Fig. 1 A). The HEK cells were cotransfected with a plasmid containing a luciferase reporter gene under the control of five <u>sis</u>-inducible elements and a plasmid encoding *Renilla* luciferase. Transfection efficiency was similar for HEK cells transfected with WT or mutant *IL23R* cDNAs, as indicated by the proportion of mCherry^+^ and IL-12Rβ1^+^ cells and the similar levels of *IL23R* and *IL12RB1* mRNAs (Fig. S1 A). We then evaluated the response to stimulation with 10 ng/ml IL-23 by assessing luciferase activity. HEK cells transfected with the mutant *IL23R* cDNAs had activity levels similar to (Q3H, A55T, R86Q, K150T, V159M, V160A, L193F, A199V, S221F, P306S, L310P, V362I, I373F, Q487H, and S559R) or lower (G300V, R381Q, G149R, and L372F) than that of cells transfected with WT *IL23R* cDNA (Fig. 1 A) or had no activity at all (C115Y, as a representative of known complete LOF variants). The allele carrying both G300V and V362I behaved like G300V, and the degree of hypomorphism could be ranked as follows, from low to high levels of luciferase activity: L372F < G300V < G149R < R381Q. STAT3 phosphorylation following IL-23 stimulation was assessed by western blotting of lysates prepared from HEK cells transfected with *IL12RB1* and either WT or mutant *IL23R* cDNAs. HEK cells transfected with the four mutated *IL23R* cDNAs (G300V, R381Q, G149R, and L372F), as identified in the luciferase assay, had lower levels of STAT3 phosphorylation following IL-23 stimulation than cells transfected with the WT *IL23R* cDNA. No STAT3 phosphorylation upon IL-23 stimulation was detected with the known LOF variant C115Y (Fig. 1 B, Table 1, and Fig. S1 B). Overall, 15 variants were found to be isomorphic, four were hypomorphic (≤50% of WT activity), and the previously reported C115Y variant (Martinez-Barricarte et al., 2018) was LOF for cellular responses to IL-23 in this overexpression setting.

**Table 1.**
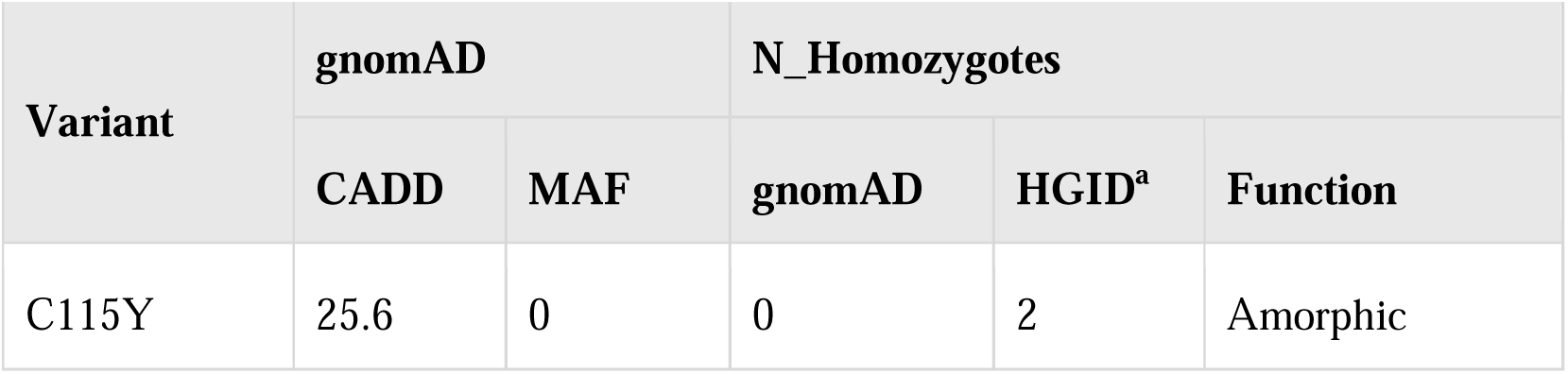
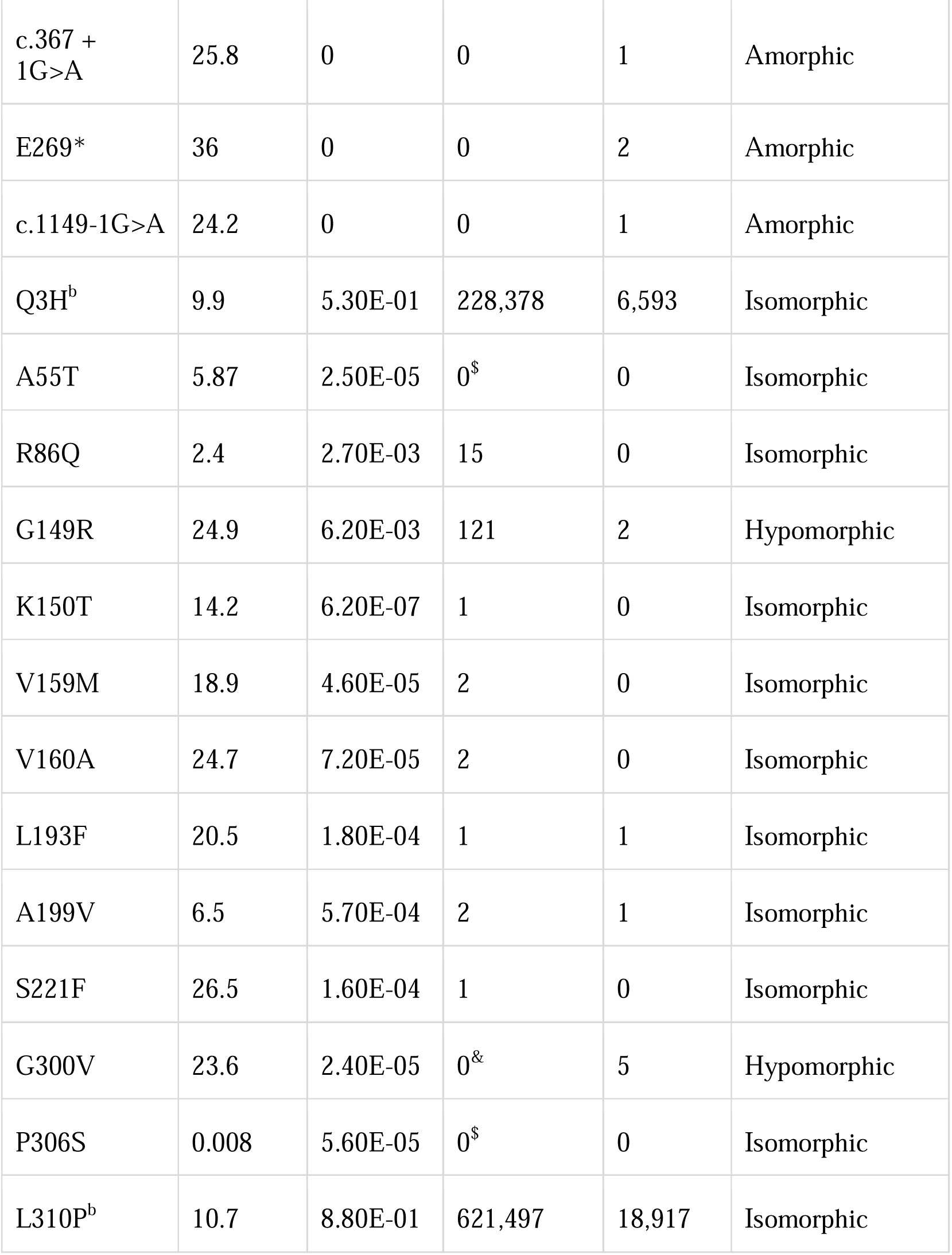
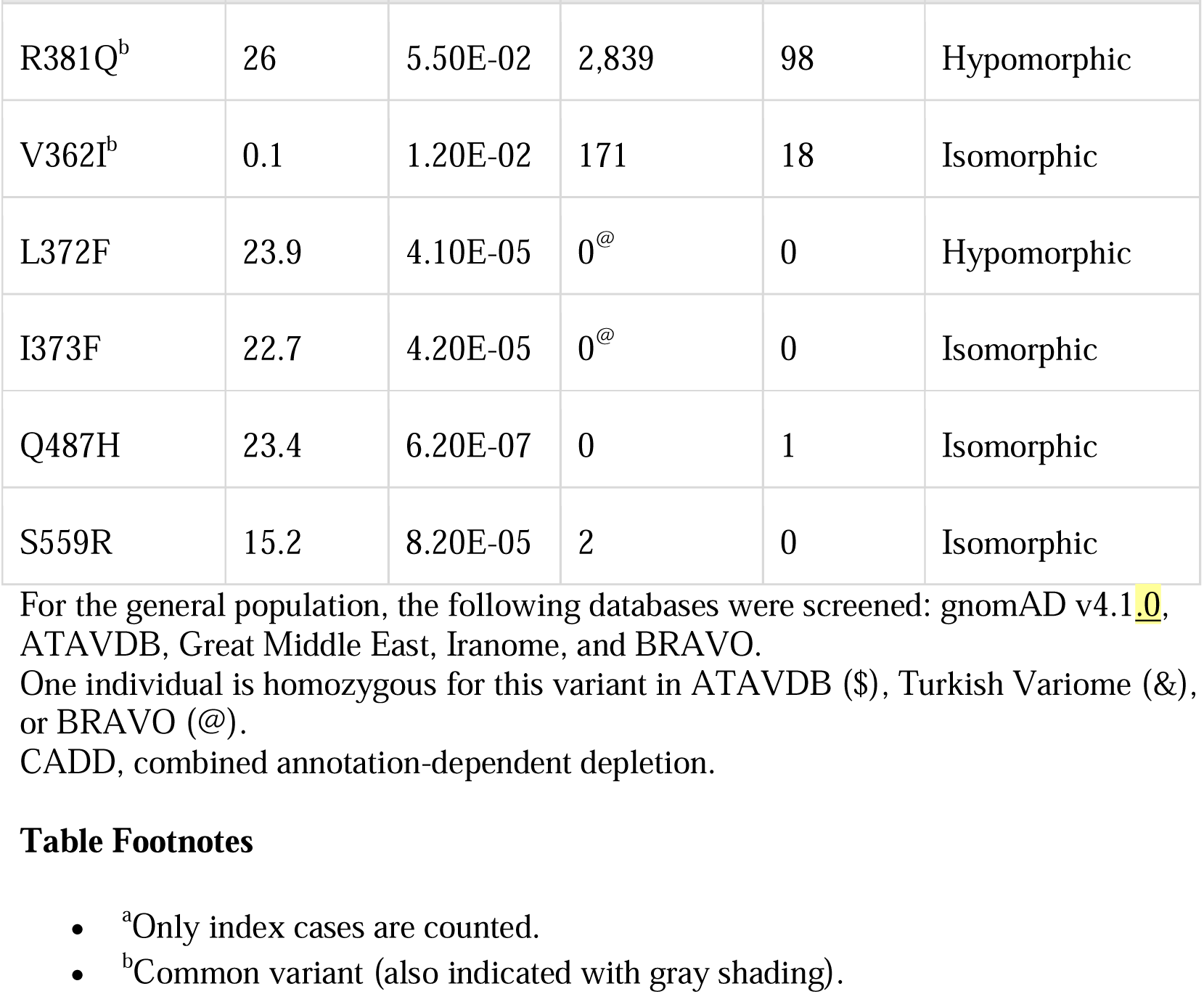
Biallelic *IL23R* coding variants in our in-house cohort of 24,046 patients and in cohorts of individuals from the general population.

| Variant | gnomAD |  | N_Homozygotes |  |  |
| --- | --- | --- | --- | --- | --- |
|  | CADD | MAF | gnomAD | HGID<sup>a</sup> | Function |
| C115Y | 25.6 | 0 | 0 | 2 | Amorphic |
|  | CADD | MAF | gnomAD | HGID<sup>a</sup> | Function |
| c.367 + 1G>A | 25.8 | 0 | 0 | 1 | Amorphic |
| E269* | 36 | 0 | 0 | 2 | Amorphic |
| c.1149-1G>A | 24.2 | 0 | 0 | 1 | Amorphic |
| Q3H<sup>b</sup> | 9.9 | 5.30E-01 | 228,378 | 6,593 | Isomorphic |
| A55T | 5.87 | 2.50E-05 | 0<sup>\$</sup> | 0 | Isomorphic |
| R86Q | 2.4 | 2.70E-03 | 15 | 0 | Isomorphic |
| G149R | 24.9 | 6.20E-03 | 121 | 2 | Hypomorphic |
| K150T | 14.2 | 6.20E-07 | 1 | 0 | Isomorphic |
| V159M | 18.9 | 4.60E-05 | 2 | 0 | Isomorphic |
| V160A | 24.7 | 7.20E-05 | 2 | 0 | Isomorphic |
| L193F | 20.5 | 1.80E-04 | 1 | 1 | Isomorphic |
| A199V | 6.5 | 5.70E-04 | 2 | 1 | Isomorphic |
| S221F | 26.5 | 1.60E-04 | 1 | 0 | Isomorphic |
| G300V | 23.6 | 2.40E-05 | 0<sup>&</sup> | 5 | Hypomorphic |
| P306S | 0.008 | 5.60E-05 | 0<sup>\$</sup> | 0 | Isomorphic |
| L310P<sup>b</sup> | 10.7 | 8.80E-01 | 621,497 | 18,917 | Isomorphic |
|  | CADD | MAF | gnomAD | HGID<sup>a</sup> | Function |
| R381Q<sup>b</sup> | 26 | 5.50E-02 | 2,839 | 98 | Hypomorphic |
| V362I<sup>b</sup> | 0.1 | 1.20E-02 | 171 | 18 | Isomorphic |
| L372F | 23.9 | 4.10E-05 | 0<sup>@</sup> | 0 | Hypomorphic |
| I373F | 22.7 | 4.20E-05 | 0<sup>@</sup> | 0 | Isomorphic |
| Q487H | 23.4 | 6.20E-07 | 0 | 1 | Isomorphic |
| S559R | 15.2 | 8.20E-05 | 2 | 0 | Isomorphic |

For the general population, the following databases were screened: gnomAD v4.1.0, ATAVDB, Great Middle East, Iranome, and BRAVO. One individual is homozygous for this variant in ATAVDB (\$), Turkish Variome (&), or BRAVO (@). CADD, combined annotation-dependent depletion.

**Figure 1:**
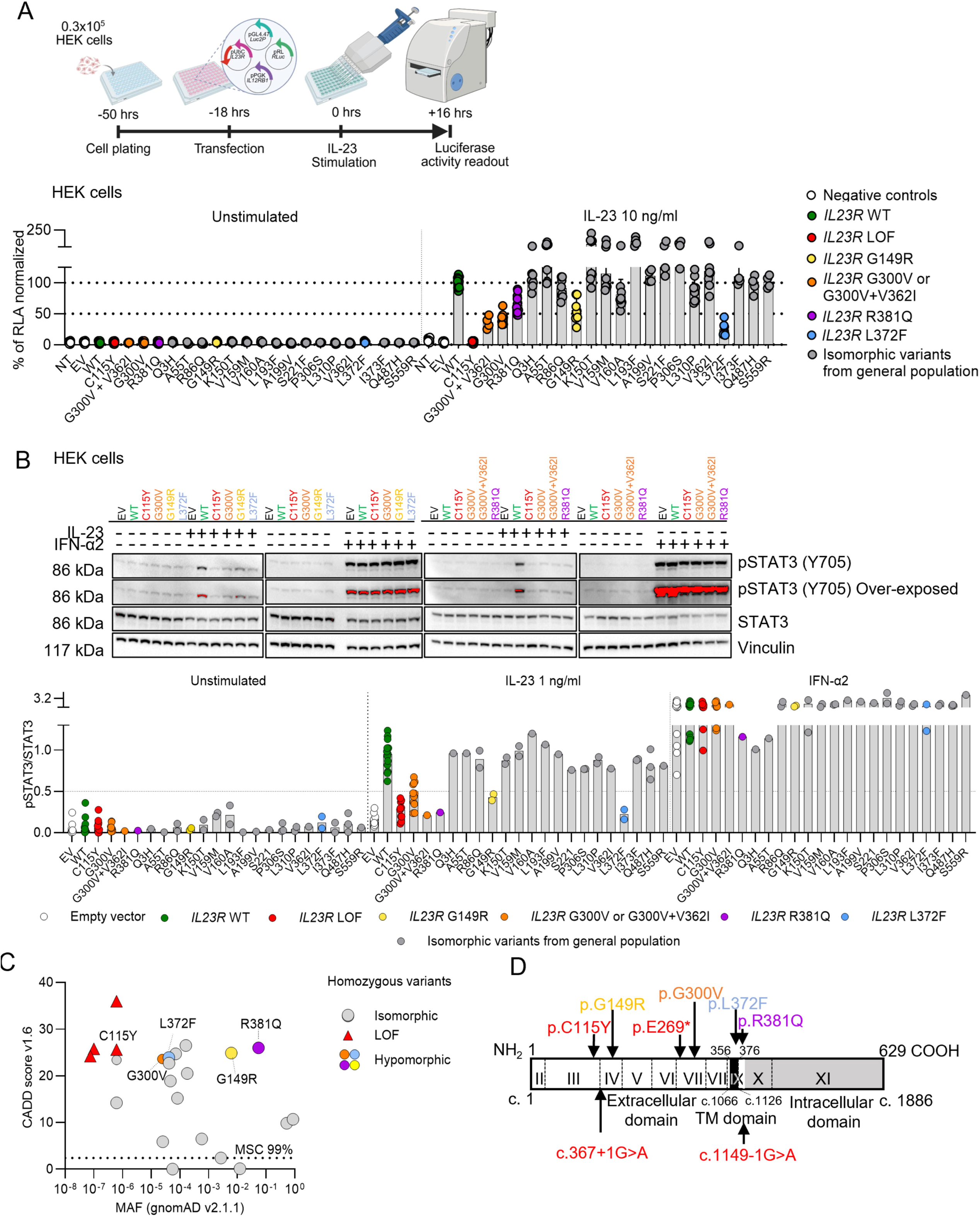
Images description Functional impact of *IL23R* variants on IL-23–dependent STAT3 activity. **(A)** Schematic representation of the experiment and STAT3 activity induced in response to IL-23 (10 ng/ml) by the *IL23R* variants used for the transient transfection of HEK cells, assessed with the luciferase assay. White dots represent negative controls (NT, not transfected and EV, empty vector); green dots, WT condition; red dots, LOF variant; gray dots, homozygous variants from the general population. Each variant was transfected at least five times, and each data point is represented by a circle. **(B)** Western blot of pSTAT3 after stimulation with IL-23 (1 ng/ml) or IFN-α (1 ng/ml) in HEK cells transiently transfected with *IL23R* variants. The graph shows pSTAT3/STAT3 band density as a percentage of that in stimulated WT conditions. At least two independent experiments were performed. The blot is duplicated in Fig. S1 B. **(C)** CADD/MAF graph for the homozygous *IL23R* LOF variants previously described (red triangle) in the MSMD cohort and the missense variants (dots) from the general population and our in-house cohort. The four hypomorphic variants are presented in different colors, whereas the isomorphic variants are shown in gray. MSC, mutation significance cutoff with a 99% CI. CADD, combined annotation-dependent depletion. **(D)** Localization of coding and non-coding LOF (red) or hypomorphic (orange, yellow, blue, and violet) *IL23R* variants across IL-23R protein domains. Source data are available for this figure: SourceData F1.

### Population genetics of *IL23R*

The consensus negative selection score (CoNeS = −0.08) and the gene damage index (GDI = 3.6) of *IL23R* are similar to those of *IL12RB1* (CoNeS = 0.45; GDI = 3.79), *IL12RB2* (CoNeS = 1.02; GDI = 4.05), and *TYK2* (CoNeS = −0.07; GDI = 7.4). These values are consistent with deleterious variants of the gene having the potential to underlie an AR inborn error of immunity (Itan et al., 2015; Rapaport et al., 2020). No biallelic pLOF *IL23R* variants were found in public databases (gnomAD v4.1.0, ATAVDB, Great Middle East, Iranome, Turkish Variome, and BRAVO). The hypomorphic G300V and L372F variants are rare in the general population (frequency in gnomAD v4.1.0 = 0.0024% and 0.004%, respectively), with the highest frequencies recorded in populations of Middle Eastern ancestry for G300V (frequency in this population in gnomAD v4.1.0 = 0.032%) and East Asian ancestry for L372F (frequency in this population in gnomAD v4.1.0 = 0.118%) (Fig. 1 C, Fig. S2 A, and Table 1). In the Turkish Variome database (Kars et al., 2021), the G300V variant was reported only once in the homozygous state, and there were no individuals homozygous for L372F. In our own database (Human Genetic of Infectious Diseases [HGID] database), L372F is absent, and G300V was reported in the homozygous state in two index individuals of European ancestry. We estimated that the most recent common ancestor carrying the G300V variant lived about 3,078 years ago (95% confidence interval [CI]: 972–11,500 years). The G149R variant is more frequent than G300V and L372F (frequency in gnomAD v4.1.0 = 0.616%), and its frequency is highest in East Asian populations (frequency in this population in gnomAD v4.1.0 = 6%), with a total of 121 homozygotes identified in gnomAD v4.1.0 (0.01%, mainly in East Asian and non-Finnish Europeans) and two unrelated individuals identified in our HGID cohort. The R381Q variant is the most frequent of the hypomorphic variants (frequency in gnomAD v4.1.0 = 5.4%), with the highest frequency observed in Amish, Ashkenazi Jewish, and non-Finnish European populations (frequency in these populations in gnomAD v4.1.0 = 10, 7.2, and 6.4% respectively). It was reported in the homozygous state in 2,839 individuals (0.4%) in the gnomAD v4.1.0 database and in 100 individuals (0.4%) in our HGID cohort (Table S1). Overall, the cumulative frequency of the hypomorphic homozygous variants G149R and R381Q in gnomAD v4.1.0 was 0.4%, ranging from 0.02% in the African population to 1.3% in the Amish population.

### The IL-23R mutants have low levels of cell surface expression and/or effects on receptor conformation potentially affecting IL-23–induced STAT3 phosphorylation

Investigations of the impact of the C115Y variant on IL-23R cell surface expression indicated a small decrease in expression in HEK cells (Lay et al., 2023). However, the lack of a functional response of C115Y IL-23R to IL-23 results predominantly from a disruption of IL-23 binding (Lay et al., 2023). We focused on the four newly discovered hypomorphic variants. The G149R and G300V variants localize to the extracellular domain, whereas the L372F and R381Q variants localize to the transmembrane and intracellular domains of IL-23R, respectively (Fig. 1 D) (Parham et al., 2002). The cell surface expression of the G300V and R381Q variants was weaker than that of WT IL-23R, at levels similar to those reported for the C115Y variant (Fig. 2, A and B) (Lay et al., 2023). By contrast, the cell-surface expression of the G149R and L372F variants was similar to that of WT IL-23R (Fig. 2, A and B). Wide-field luminescence imaging demonstrated receptor expression in the form of luminescence originating from the N-terminal NLuc tag fused to the variant or WT IL-23R following the addition of the NLuc substrate furimazine (Fig. 2 B). We previously performed NanoLuciferase bioluminescence resonance energy transfer (NanoBRET) assays between N-terminal NanoLuciferase-tagged WT IL-23R or IL-12Rβ1 and a TAMRA-labeled IL-23 (Lay et al., 2022). These experiments demonstrated a 66-fold increase in the affinity of IL-23–TAMRA for the heterodimeric complex composed of NLuc–IL-23R and unlabeled IL-12Rβ1 (K_D_ = 27.0 pM) relative to NLuc–IL-23R or NLuc–IL-12Rβ1 expressed individually (K_D_ = 222.2 nM or 30.1 nM, respectively). Here, we performed comparable experiments with the four IL-23R variants, each tagged at the N terminus with NanoLuc and used for cotransfection with IL-12Rβ1. All four IL-23R variants had binding affinities similar to that of WT IL-23R (Fig. 2 C and Fig. S2 B). Importantly, these K_D_ values were substantially lower than the concentrations required for binding to IL-23R or IL-12Rβ1 expressed individually (Lay et al., 2022), suggesting that the measurements were unlikely to be confounded by IL-23–TAMRA binding to non-heterodimeric receptor complexes at the higher ligand concentrations used. However, the lower maximum NanoBRET signal observed for G300V and G149R and, to a lesser extent, R381Q and L372F probably reflect changes in the conformation of the receptor altering its orientation or the distance between the NanoLuciferase donor tag and the TAMRA fluorophore acceptor, resulting in a lower efficiency of energy transfer for these mutant receptors. None of the variants significantly altered the ability of IL-23R and IL-12Rβ1 to form dimers in the presence or absence of 5 nM IL-23, although the lower levels of IL-23R at the surface reduced overall levels of dimer formation in both instances (Fig. S2 C). These data suggest that the weaker pSTAT3 responses observed with the G300V, R381Q, G149R, and L372F variants (Fig. 2, A–C; and Fig. S2, B and C) are not due to a lower affinity for IL-23, but instead reflect the lower levels of cell surface expression for some of these IL-23R variants (R381Q and G300V) and/or conformational changes affecting the efficacy of the agonist with respect to STAT3 phosphorylation.

**Figure 2:**
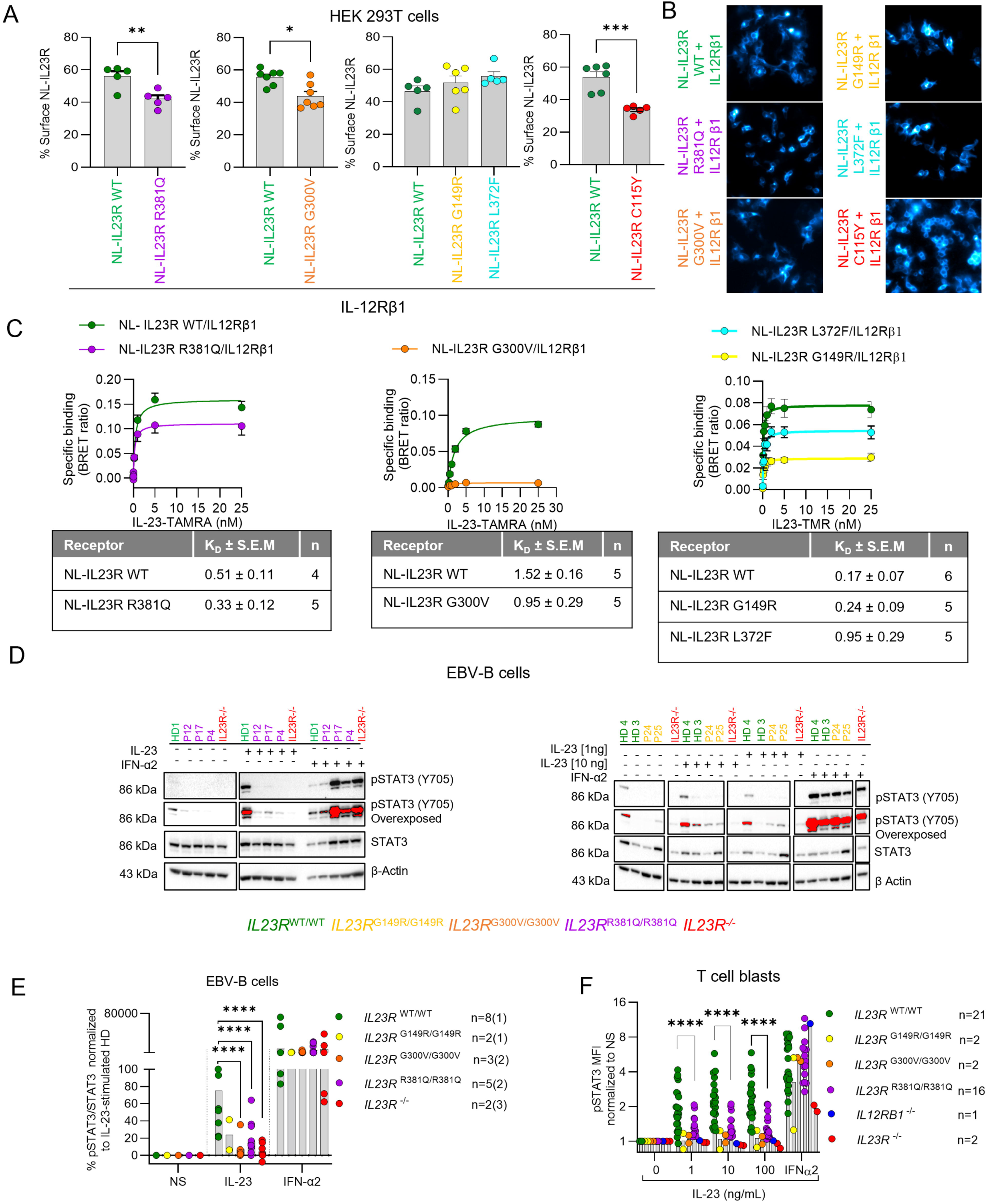
Impaired expression and/or conformation of IL-23R and responses to IL-23 in patient cell lines. **(A)** Emitted luminescence measured in the presence of the NanoLuc inhibitor is expressed as a percentage of that measured in the absence of the inhibitor (100%) in HEK cells transiently transfected with N-terminal NanoLuciferase-tagged versions of the WT or variants of the IL-23 receptor (NLuc IL-23R, NLuc IL-23R R381Q, NLuc IL-23R G300V, NLuc IL-23R G149R, NLuc IL-23R L372F, or NLuc IL-23R C115Y) and IL-12Rβ1 (4:1 ratio). The data shown are the mean ± SEM from five to seven independent experiments conducted in triplicate wells. Statistical significance was determined in a paired *t* test (***P < 0.001, **P < 0.01 and *P < 0.05). Data for NLuc IL-23R C115Y were obtained in a previous study (Lay et al., 2023). **(B)** HEK cells transiently transfected as described in A were imaged with an Olympus LuminoView 200 wide-field luminescence microscope. Representative luminescence images show the signal originating from the N-terminal NLuc tag of each IL-23R variant following the addition of furimazine (final dilution 1:400). Data from three independent experiments are shown. The scale bar represents 50 μm. Luminescence images were acquired with a 60× NA1.42 oil immersion objective, a 0.5× tube lens, and a C9100-23B IMAGE EMX2 camera (Hamamatsu, Japan), with an exposure time of 20 s (gain of 25). **(C)** HEK cells transiently transfected as in A were treated with various concentrations of fluorescently labeled IL-23 (IL-23–TAMRA) in the presence or absence of unlabeled IL-23. The NanoLuciferase substrate furimazine was then added (final concentration: 7.7 μM), and emitted luminescence and fluorescence were simultaneously detected with a BMG PHERAstar FS. BRET ratios were calculated by subtracting fluorescence from luminescence. The specific binding of IL-23–TAMRA was calculated by subtracting BRET ratios determined in the presence of unlabeled IL-23 (nonspecific binding) from those obtained in its absence. The data shown are the mean ± SEM (*n* = 4–5 independent experiments performed in triplicate wells for NLuc IL-23R WT versus NLuc IL-23R R381Q; *n* = 5 independent experiments performed in triplicate for NLuc IL-23R WT versus NLuc IL-23R G300V; *n* = 5 independent experiments performed in duplicate for NLuc IL-23R WT versus NLuc IL-23R G149R or NLuc IL23R L372F). **(D)** Representative results for pSTAT3 detection by western blotting after IL-23 stimulation (10 ng/ml and/or 1 ng/ml) in EBV–B cell lines from healthy controls (HD: green), *IL23R*^R381Q/R381Q^ (P12, P17, and P4: violet), *IL23R*^G300V/G300V^ (P18, P19, and P20: orange), *IL23R*^G149R/G149R^ (P24 and P25: yellow), and *IL23R*^LOF/LOF^ (*IL-23R*^−/−^: red) patients. **(E)** Representative graph of multiple independent western blot experiments showing pSTAT3/STAT3 band density. The number of cells per genotype is indicated in the figure (*n* =). If there were several measurements on the same EBV–B cells, this is indicated in parentheses. Statistical significance was assessed in unpaired Mann–Whitney *U* tests comparing each variant with HD or WT as appropriate (****P < 0.0001). **(F)** MFI of pSTAT3 normalized against non-stimulated T cell blasts from healthy controls (green dots), *IL23R*^G149R/G149R^ (yellow dots), *IL23R*^G300V/G300V^ (orange dots), *IL23R*^R381Q/R381Q^ (purple dots), and IL-12Rβ1– and IL-23R–deficient patients (blue and red dots, respectively) in the presence or absence of various concentrations of IL-23. IFNα2 stimulation (1 ng/ml) was used as a control. Statistical significance was assessed in unpaired Mann–Whitney’s *U* tests comparing each variant with HD or WT as appropriate (****P < 0.0001). Source data are available for this figure: SourceData F2.

### Impaired IL-23 signaling in lymphoid cell lines derived from patients homozygous for the *IL23R* variants

We then used patient-derived cell lines to capture the impact of the full *IL23R* locus genotypes of the patients in the context of their own genome. We obtained EBV-immortalized B (EBV-B) cells from eight healthy controls, three patients homozygous for G300V, five patients homozygous for R381Q, and two patients homozygous for G149R (*IL23R*^WT/WT^, *IL23R*^G300V/G300V^, *IL23R*^R381Q/R381Q^, and *IL23R*^G149R/G149R^ EBV-B cells, respectively). We also included cells from two patients with complete IL-23R deficiency (C115Y: *IL23R*^−/−^) in the analysis. However, we were unable to derive EBV-B cells from an individual homozygous for L372F, as no such individuals are present in our in-house cohort (>24,000 individuals). These EBV-B cells were left unstimulated or were stimulated with IL-23 or with IFN-α2a as a positive control. EBV-B cells from healthy controls (*IL23R*^WT/WT^) responded to IL-23 by STAT3 phosphorylation (Fig. 2, D and E; and Fig. S2 D). In *IL23R*^R381Q/R381Q^, *IL23R*^G149R/G149R^, and *IL23R*^G300V/G300V^ EBV-B cells, STAT3 phosphorylation in response to IL-23 was impaired but not abolished. By contrast, STAT3 was phosphorylated in response to IFN-α2a in all cells (Fig. S2 D). In parallel, we assessed the functionality of *IL23R* variants in T cell blasts derived from 21 healthy controls, 2 *IL23R*^G300V/G300V^, 16 *IL23R*^R381Q/R381Q^, 2 *IL23R*^G149R/G149R^, 2 *IL23R*^−/−^, and 1 *IL12RB1*^−/−^ patient. T cell blasts were left unstimulated or were stimulated with IL-23 or IFN-α2a and STAT3 phosphorylation was monitored by intracellular flow cytometry. T cell blasts from *IL23R*^WT/WT^ individuals responded to IL-23 by phosphorylating STAT3, whereas no such phosphorylation was observed in cells from *IL23R*G300V/G300V, *IL23R*^R381Q/R381Q^, *IL23R*^G149R/G149R^, *IL23R*^−/−^, and *IL12R*β*1*^−/−^ patients (Fig. 2 F), as previously shown for *IL23R*^R381Q/R381Q^ (Gerbaux et al., 2025). All T cell blasts responded similarly to IFN-α2a. Thus, IL-23 signaling was impaired but not abolished in lymphoid cells derived from patients homozygous for the G149R, G300V, or R381Q *IL23R* variants.

### Impaired IL-23–mediated *IFNG* mRNA induction in *IL23R*^G300V/G300V^ and *IL23R*^R381Q/R381Q^ leukocytes

Human IFN-γ is essential for antimycobacterial immunity, as most of the known genetic etiologies of MSMD or TB are associated with impaired IFN-γ activity (Boisson-Dupuis et al., 2024; Casanova et al., 2024). We therefore studied the impact of the three *IL23R* hypomorphic genotypes on IL-23–dependent IFN-γ production. We first compared the distributions of peripheral blood mononuclear cells (PBMCs) from healthy controls, nine *IL23R*^R381Q/R381Q^ patients, one *IL23R*^G300V/G300V^ patient, one *IL23R*^G149R/G149R^ patient, one *IL12RB1*^−/−^ patient, and two *IL23R*^−/−^ patients by CyTOF with two antibody panels (Fig. S3, A–D). Like patients with AR complete IL-23R deficiency, all the patients studied here had normal counts and frequencies of myeloid and lymphoid subsets, including normal frequencies of helper T cells, natural killer (NK) cells, and MAIT and γδ T cells, which produce large amounts of IFN-γ upon IL-23 stimulation (Martinez-Barricarte et al., 2018; Philippot et al., 2023) (Fig. S3, A–D). We also performed baseline single-cell RNA sequencing (scRNAseq) on two *IL23R*^G300V/G300V^, three *IL23R*^R381Q/R381Q^ patients, three *TYK2*^P1104A/P1104A^, one *IL23R*^G149R/G149R^, three *IL23R*^−/−^, three *IL12RB1*^−/−^ patients, and 11 healthy controls. Clustering analysis revealed comparable numbers and proportions of the 22 discrete transcriptionally defined leukocyte subsets detected in healthy controls and the patients (Fig. 3 A and Fig. S4, A and B), consistent with data derived from flow cytometric immunophenotyping. Gene set enrichment analysis (GSEA) revealed a downregulation of genes regulated by IFN-γ in the classical monocytes of all patients homozygous for hypomorphic *IL23R* alleles (G149R, G300V, and R381Q), as previously described for IL-23R^−/−^, IL-12Rβ1^−/−^, and TYK2 P1104A deficiency (Fig. 3 B) (Martinez-Barricarte et al., 2018; Ogishi et al., 2022; Philippot et al., 2023). We hypothesized that homozygosity for a hypomorphic *IL23R* variant might affect *IFNG* mRNA induction following the stimulation of patient leukocytes with IL-23 stimulation. We tested this hypothesis with two *IL23R*^G300V/G300V^, one *IL23R*^G149R/G149R^, three *IL23R*^R381Q/R381Q^, two *TYK2*^P1104A/P1104A^, two *IL23R*^−/−^, and one *IL12RB1*^−/−^ patient, and nine healthy controls, by performing scRNAseq on PBMCs incubated with or without IL-23 for 6 h (Fig. 3, C and D; and Fig. S4, C and D). As previously reported in patients with complete IL-23R or IL-12Rβ1 deficiency, NK, MAIT, and Vδ2^+^ γδ T cells homozygous for the *IL23R* G149R, *IL23R* G300V, or *TYK2* P1104A variant displayed similar impairments of *IFNG* mRNA induction upon IL-23 stimulation (Ogishi et al., 2022; Philippot et al., 2023). By contrast, a milder impairment was observed in cells homozygous for the *IL23R* R381Q variant, consistent with the degree of hypomorphism in the luciferase assay (Fig. 1 A and Fig. 3 D). Overall, IL-23–mediated *IFNG* mRNA induction was impaired following IL-23 stimulation in MAIT, NK, and Vδ2^+^ γδ T cells from patients homozygous for hypomorphic IL-23R G300V, G149R, and R381Q variants and *TYK2*^P1104A/P1104A^ individuals.

**Figure 3:**
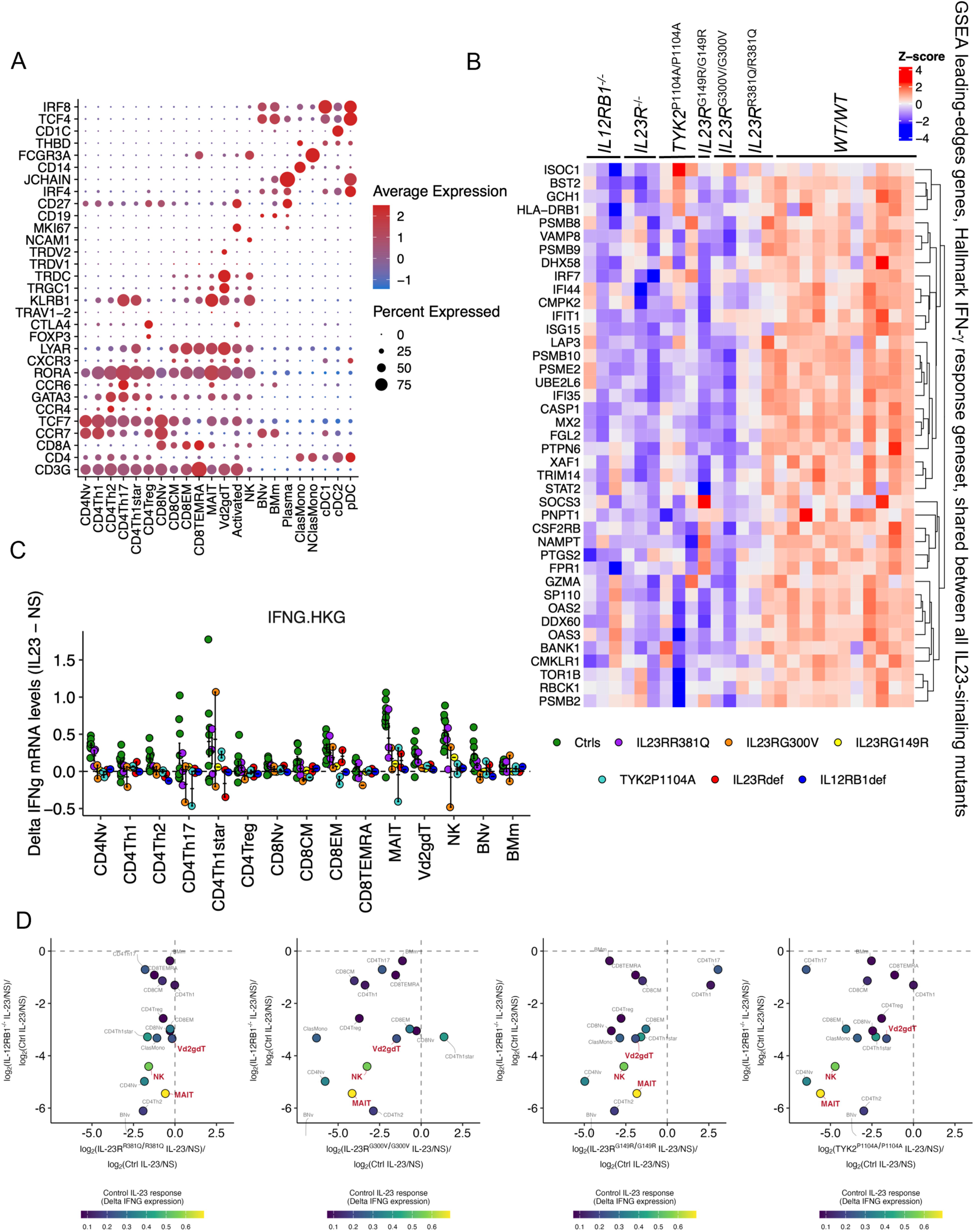
Impaired IL-23-mediated *IFNG* mRNA induction in patients homozygous for hypomorphic *IL23R* variants. **(A)** Dotplot of the expression of lineage- and state-specific marker genes across predicted immune cell types. **(B)** Heatmap analysis of the GSEA leading-edge genes for the Hallmark IFN-γ response gene set common to IL-23R–deficient and IL-12Rβ1–deficient classic monocytes. Normalized *Z*-transformed pseudobulk read counts are shown. Three *IL12RB1*^−/−^, three *IL23R*^−/−^, three *TYK2*^P1104A/P1104A^, one *IL23R*^G149R/G149R^, two *IL23R*^G300V/G300V^, three *IL23R*^R381Q/R381Q^ patients and 12 healthy controls were included. **(C)** The fold-change in *IFNG* mRNA levels following IL-23 stimulation in leukocytes from three *IL23R*^R381Q/R381Q^, two *IL23R*^G300V/G300V^, one *IL23R*^G149R/G149R^ and two *TYK2*^P1104A/P1104A^, one *IL12RB1*^−/−^ and two *IL23R*^−/−^ patients, and nine healthy controls was normalized against housekeeping genes across immune cells. Bulk RNAseq pseudobulk expression counts for *IFNG* were normalized by this factor to obtain IFNG.HKG values. The delta score is the difference in normalized expression between IL-23 stimulation and non-stimulation (NS) conditions (IL-23 − NS), reflecting the magnitude of gene induction upon stimulation. **(D)** Two-dimensional plots of IL-23–induced delta *IFNG* expression. The fold-change difference in *IFNG* mRNA levels following IL-23 stimulation in leukocytes from *IL23R*^R381Q/R381Q^, *IL23R*^G300V/G300V^, *IL23R*^G149R/G149R^, and *TYK2*^P1104A/P1104A^ patients relative to controls is shown on the x-axis. The y-axis shows the same parameter for IL23R^−/−^ patients as a comparison. The color of the circles indicates the median fold-change difference (IL-23 versus NS) in normalized IFNG mRNA levels in controls for the corresponding subsets. MAIT, NK, and Vδ2^+^ γδT cells are highlighted.

### Impaired *ex vivo* IL-23–mediated induction of IFN-**γ** by innate lymphoid and innate-like T cells homozygous for *IL23R* hypomorphic variants

We then used spectral flow cytometry to assess intracellular IL-23–dependent IFN-γ production by specific lymphocyte subsets: CD4^+^, CD8^+^, NK (bright/dim) and MAIT, and γδ T (Vδ1^+^ and Vδ2^+^) cells. IFN-γ induction in response to IL-23 is barely detectable in CD4^+^, CD8^+^, and Vδ1^+^ γδ T cells, probably because they express IL-23R only weakly, if at all, in the basal state (Camard et al., 2025). As previously described (Bohlen et al., 2023; Philippot et al., 2023), stimulation with IL-18 and high doses of IL-23 stimulation induced a synergistic increase in the frequency of IFN-γ–expressing MAIT, Vδ2^+^ γδ T, and NK cells relative to stimulation with IL-18 only in healthy controls. No such effect was observed with *IL23R*^G300V/G300V^, *IL23R*^G149R/G149R^, and *IL12RB1*^−/−^ cells, and a less pronounced defect was observed in *IL23R*^R381Q/R381Q^ and *TYK2*^P1104A/P1104A^ cells (Fig. S5, A–C). However, a clear defect was observed in IFN-γ^+^ cells stimulated with a lower dose of IL-23 (1 ng/ml) for MAIT, Vδ2^+^ γδ T, and NK cells from *IL23R*^R381Q/R381Q^, *IL23R*^G300V/G300V^, *IL23R*^G149R/G149R^, and *TYK2*^P1104A/P1104A^ individuals, as also observed in *IL12RB1*^−/−^ and *IL23R*^−/−^ cells, relative to cells from healthy individuals (Fig. 4, A–C). To establish that this does not reflect a generalized inability of the cells to produce cytokines *in vitro*, the production of IFN-γ and TNF following stimulation with PMA ionomycin was also assessed and was found similar for all genotypes (Fig. 4). After infection with BCG, intracellular IFN-γ induction was impaired in MAIT, Vδ2^+^ γδ T, and NK from *IL23R*^G300V/G300V^, *IL23R*^G149R/G149R^, and *IL23R*^R381Q/R381Q^ patients and from a *TYK2*^P1104A/P1104A^ individual (Fig. 5, A–C). These data suggest that homozygosity for any of the three hypomorphic IL-23R variants (G149R, G300V, and R381Q) impairs IL-23–dependent IFN-γ production *ex vivo* but does not totally abolish it, as reported for *TYK2*^P1104A/P1104A^. In addition, the IL-17A induction induced by PMA ionomycin stimulation on lymphocytes was intermediate between that of healthy individuals and *IL12RB1*^−/−^ and *IL23R*^−/−^ patients (Fig. 5 D), consistent with the absence of chronic CMC in these patients. The impairment of IL-23–dependent IFN-γ production mostly affected MAIT, Vδ2^+^ γδ T, and NK cells, resulting in a predisposition to TB.

**Figure 4:**
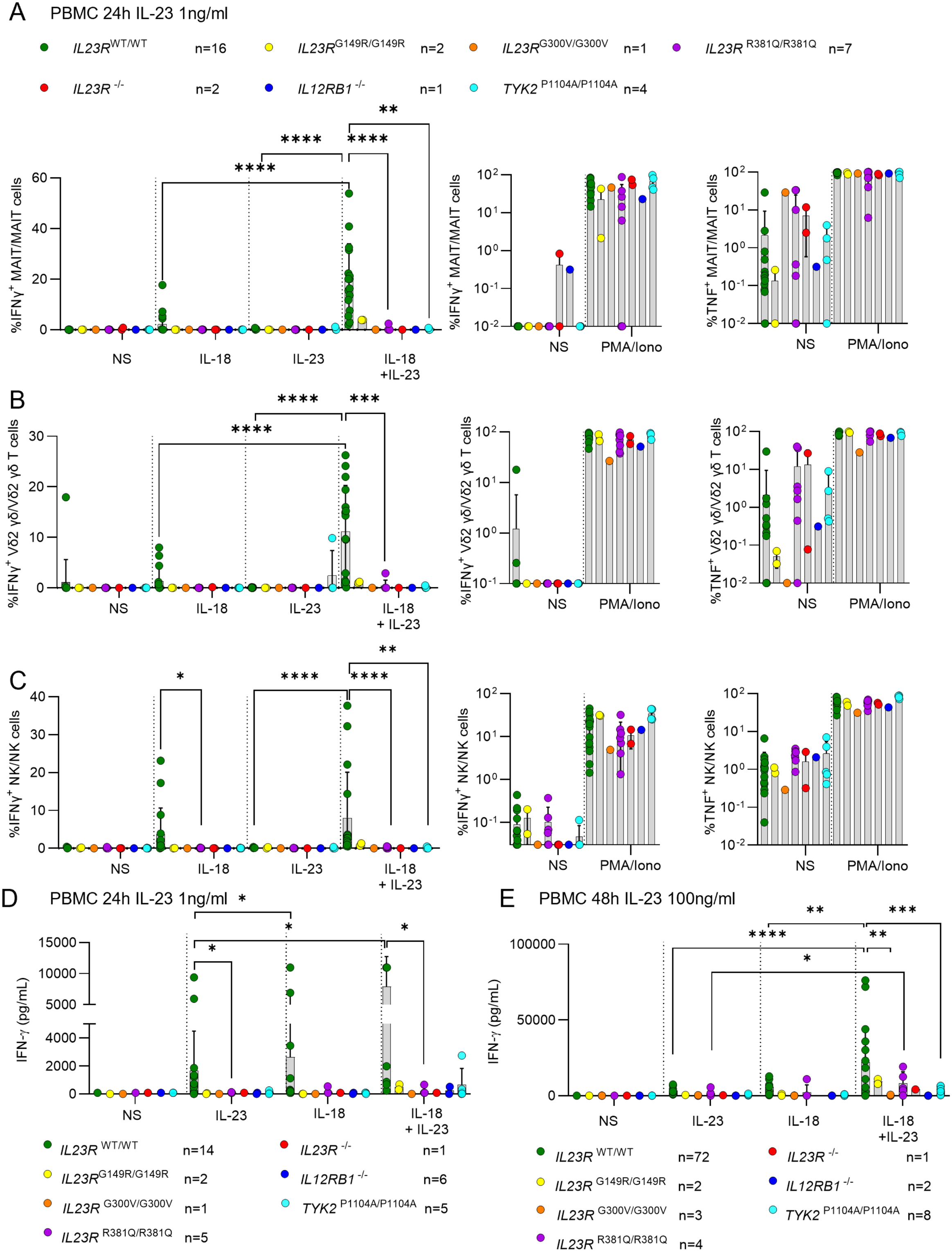
Impaired IL-23-dependent induction of IFN-γ in MAIT, Vδ2^+^ γδ T, and NK cells. **(A–C)** Percent intracellular IFN-γ^+^ induction was monitored by spectral flow cytometry in MAIT (A), Vδ2^+^ γδ T cells (B), and NK cells (C) after stimulation with IL-18 (200 ng/ml) and IL-23 (1 ng/ml), alone or in combination for 24 h, or with PMA/ionomycin stimulation for 6 h for frozen PBMCs. The percentages of IFN-γ^+^ and TNF^+^ cells were monitored as a control. 16 healthy controls, 2 *IL23R*^G149R/G149R^, 1 *IL23R*^G300V/G300V^, 7 *IL23R*^R381Q/R381Q^, 2 *IL23R*^−/−^, 1 *IL12RB1*^−/−^, and 4 *TYK2*^P1104A/P1104A^ patients were included. The statistical significance of differences was assessed in unpaired Mann–Whitney *U* tests for comparisons of each variant with HD or WT as appropriate. *P < 0.05, **P < 0.01, ***P < 0.001, ****P < 0.0001. **(D)** IFN-γ levels in the supernatant were assessed after the stimulation of thawed PBMCs of the indicated genotypes for 24 h with IL-23 (1 ng/ml), IL-18 (200 ng/ml), or both. 14 healthy controls, 2 *IL23R*^G149R/G149R^, 1 *IL23R*^G300V/G300V^, 5 *IL23R*^R381Q/R381Q^, 1 *IL23R*^−/−^, 6 *IL12RB1*^−/−^, and 5 *TYK2*^P1104A/P1104A^ patients were included. Nonparametric Mann–Whitney *U* tests were used for analysis, with *P < 0.05. **(E)** IFN-γ levels in the supernatant were assessed after the stimulation of thawed PBMCs of the indicated genotypes for 48 h with IL-23 (100 ng/ml), IL-18 (25 ng/ml), or both. 17 healthy controls, 2 *IL23R*^G149R/G149R^, 3 *IL23R*^G300V/G300V^, 4 *IL23R*^R381Q/R381Q^, 1 *IL23R*^−/−^, 2 *IL12RB1*^−/−^, and 8 *TYK2*^P1104A/P1104A^ patients were included. Nonparametric Mann–Whitney *U* tests were used for analysis, with *P < 0.05, **P < 0.01, ***P < 0.001, and ****P < 0.0001.

**Figure 5:**
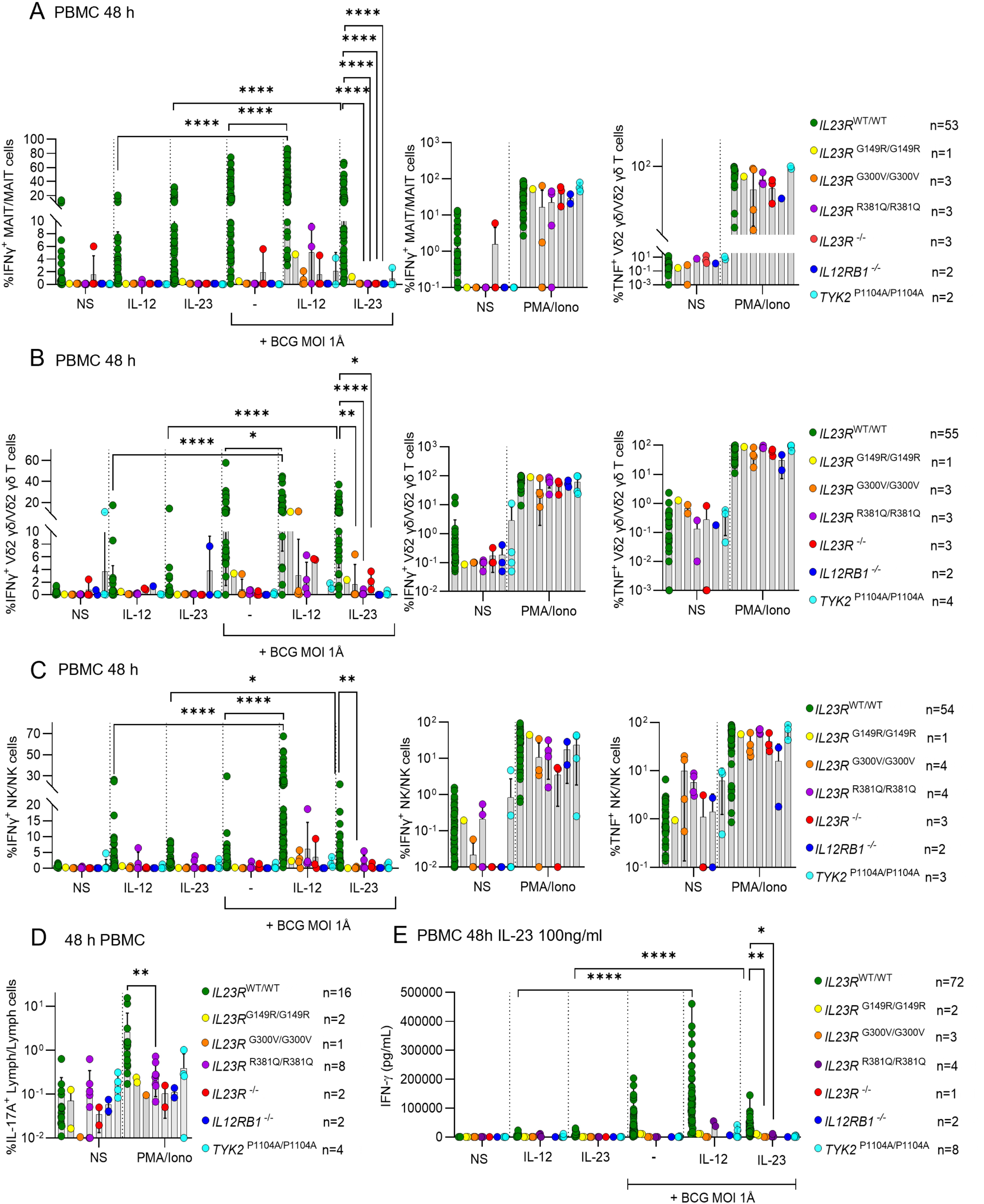
Impaired IFN-γ induction after IL-23 stimulation and mycobacterial infection. **(A–C)** Freshly thawed PBMCs of the indicated genotypes were stimulated with IL-23 (100 ng/ml) or IL-12 (5 ng/ml) in the presence or absence of BCG for 48 h or with PMA + ionomycin for 6 h. The percent IFN-γ^+^ cells was assessed: (A) among MAIT cells, (B) among Vδ2^+^ γδ T cells, (C) and among NK cells. The percentages of IFN-γ^+^ and TNF^+^ cells after stimulation with PMA + ionomycin were evaluated as controls. The number of individuals tested per genotype is indicated in each figure. Nonparametric Mann–Whitney *U* tests were used for analysis, with *P < 0.05, **P < 0.01 and ****P < 0.0001. **(D)** IL-17A induction in total lymphocytes after the stimulation with PMA of PBMCs of the indicated genotypes. The number of individuals tested per genotype is indicated in each figure. Nonparametric Mann–Whitney *U* tests were used for analysis, with **P < 0.01. **(E)** IFN-γ levels in the supernatant were assessed after the stimulation of PBMCs of the indicated genotypes for 48 h with IL-12 (5 ng/ml) or IL-23 (100 ng/ml) in the presence or absence of live *Mycobacterium bovis*–BCG (MOI 1) or PMA/ionomycin. The number of individuals tested per genotype is indicated in each figure. Nonparametric Mann–Whitney *U* tests were used for analysis, with *P < 0.05, **P < 0.01, and ****P < 0.0001.

### Homozygosity for *IL23R* hypomorphic variants impairs the *ex vivo* IL-23–mediated production of IFN-**γ**

Given the role of IL-23 in inducing IFN-γ secretion, we performed *ex vivo* experiments to test the hypothesis that homozygosity for the *IL23R* G300V, G149R, and R381Q variants impairs IL-23–dependent IFN-γ secretion. Homozygosity for *IL23R* L372F would be expected to have a similar effect, but this hypothesis could not be tested due to the absence of individuals carrying this genotype. We tested IL-23 responses either alone (at low or high concentration) or in combination with IL-1β, IL-18, or BCG in PBMCs from healthy individuals, individuals with *IL23R*^G300VG300V^, *IL23R*^R381Q/R381Q^, or *IL23R*^G149R/G149R^ variants, and in IL-23R– and IL-12Rβ1–deficient patients, with IFN-γ concentrations in the supernatant determined after 48 h (Fig. 5 E and Fig. S5 F). PBMCs from individuals with *IL23R*^G300V/G300V^, *IL23R*^G149R/G149R^, or *IL23R*^R381Q/R381Q^ genotypes produced significantly less IFN-γ upon IL-23 stimulation, with or without IL-1β, than healthy controls, with a more pronounced decrease observed in patients with complete IL-23R or IL-12Rβ1 deficiency (Fig. S5 D). A similar pattern was observed when PBMCs were stimulated with IL-23 (at low and high doses) in the presence of IL-18 (Fig. 4, D and E; and Fig. S5, E and F). Impaired IFN-γ production by *IL23R*^R381Q/R381Q^, *IL23R*^G300V/G300V^, *IL23R*^G149R/G149R^, and *TYK2*^P1104A/P1104A^ PBMCs relative to healthy controls was also observed after stimulation with BCG and IL-23 stimulation (Fig. 5 E). As a control, the induction of TNF and IFN-γ by PMA ionomycin was similar for all genotypes. Thus, IL-23–mediated IFN-γ production is impaired in *IL23R*^R381Q/R381Q^, *IL23R*^G149R/G149R^, *IL23R*^G300V/G300V^, and *TYK2*^P1104A/P1104A^ primary mononuclear cells.

### Enrichment in homozygosity for hypomorphic *IL23R* among patients with TB

The G300V rare variant was present in the homozygous state in five individuals from two unrelated Turkish families in our TB cohort (kinship coefficient of zero) (Fig. S2 E). Four of these individuals suffered from TB, none had MSMD, and the fifth remained asymptomatic. The R381Q variant is present in the homozygous state in 98 individuals from the HGID cohort: seven had TB (four North Africans, one European, one person of Middle Eastern ancestry, and one Latin American), two had MSMD, and the remaining 89 were either healthy or suffered from other infections (Table S1). Two individuals from the HGID cohort (one of Middle Eastern ancestry and one Asian) were homozygous for the G149R allele, but neither had TB or MSMD (Table S1). There were no homozygotes for L372F in our entire in-house HGID database. We first focused on the variant with the strongest hypomorphic effect, G300V, comparing the frequency of homozygosity for this variant between 1,874 TB patients of diverse ethnic origins and 20,883 individuals from the in-house HGID database (“control cohort”) who were either healthy or suffering from non-intramacrophagic diseases. The proportion of G300V homozygotes was 2.1 × 10^−3^ in the TB cohort and 4.8 × 10^−5^ in the control cohort, giving an adjusted odds ratio (OR) for TB of OR [95% CI] = 26.5 [4.2–167.0]; P value = 4.9 × 10^−4^ on logistic regression (see Materials and methods; Table S2). We then performed an activity-weighted burden test (see Materials and methods, with the weight inversely proportional to the residual activity of the variant), which demonstrated a significant enrichment in homozygosity for hypomorphic *IL23R* variants (G300V, R381Q, G149R, or L372F) among TB cases (P = 3.8 × 10^−2^) (Table S2). This enrichment was even stronger among individuals with inferred “Middle Eastern” ancestry (see Materials and methods section) (*N* cases = 404 and *N* controls = 1,871), in whom R381Q is more frequent than in other populations (frequency in the gnomAD v4 Middle Eastern population = 5.1%) (P = 9.7 × 10^−5^). By contrast, no significant enrichment in homozygosity for G300V, R381Q, G149R, or L372F was detected in cohorts of MSMD or CMC patients, as suggested in other studies (Gerbaux et al., 2025) (Table S2), and no enrichment in heterozygosity for these variants was detected in any of the cohorts, suggesting that these variants implicated in TB display recessive inheritance. As a negative control, we performed the same test on the 15 variants identified as isomorphic in the luciferase assay; we found no enrichment in these variants in the TB cohort (P = 0.48). It should be noted that the low to moderate prevalence of TB in some regions of the world (e.g., Europe and the Middle East) implies that most of the controls living in these regions included in this analysis are unlikely to have been exposed to and infected with *Mycobacterium tuberculosis.* The calculated enrichment in homozygosity for hypomorphic variants among TB patients for G300V, R381Q, G149R, or L372F is, therefore, probably an underestimation of the true risk of TB in homozygous individuals upon infection. Our data suggests that homozygosity for the G300V, R381Q, G149R, or L372F variants confers a predisposition to TB.

## Discussion

We identified rare (G149R, G300V, and L372F) and common (R381Q) *IL23R* variants that were hypomorphic. Homozygosity for each of these variants causes partial, rather than complete, IL-23R deficiency, with IL-23–dependent STAT3 phosphorylation impaired but not abolished. The underlying mechanism involves low levels of surface expression and/or conformational changes, with no effect on affinity for IL-23. Cells from individuals homozygous for the *IL23R* G300V, G149R, or R381Q variant display impaired IL-23–dependent IFN-γ production. This is probably also true for *IL23R*^L372F/L372F^, but we were unable to test cells from an individual with this genotype due to its rarity. A higher proportion of TB patients homozygous for G300V than of those homozygous for R381Q had extrapulmonary disease and an earlier disease onset. However, these observations are based on a limited number of individuals. The defect affects principally the innate-like (MAIT and Vδ2^+^ γδ T) and innate (NK) lymphocyte subsets, which are normally the cells with the best response to IL-23 stimulation (Hunter, 2005; Martinez-Barricarte et al., 2018; Ogishi et al., 2022; Philippot et al., 2023). The cellular phenotypes of individuals with the rare *IL23R*^G300V/G300V^ and *IL23R*^G149R/G149R^ genotypes or the common *IL23R*^R381Q/R381Q^ genotype resemble that of *TYK2*^P1104A/P1104A^ patients (Boisson-Dupuis et al., 2018; Ogishi et al., 2022). The residual IL-23–dependent IFN-γ production in patients homozygous for the *IL23R* G300V, G149R, and R381Q variants is sufficient to protect most of these individuals against weakly virulent mycobacteria (BCG and EM), contrasting with patients with a complete form of recessive IL-23R deficiency (Martinez-Barricarte et al., 2018; Philippot et al., 2023; Staels et al., 2022). Like *TYK2*^P1104A/P1104A^ (Boisson-Dupuis et al., 2018; Ogishi et al., 2022), *IL23R*^R381Q/R381Q^ has a very low penetrance for MSMD. By contrast, the residual IL-23–dependent IFN-γ production is not sufficient to protect homozygotes against *M. tuberculosis,* which is at least 1,000 times more virulent than BCG and EM (Boisson-Dupuis et al., 2011; Fieschi et al., 2003). Consistently, no enrichment in these rare and common *IL23R* variants was detected in the MSMD cohort. There was also no enrichment in homozygosity for these variants in the CMC cohort, consistent with the low penetrance of CMC in patients with complete IL-23R deficiency (only a third of IL-23R^−/−^ patients had CMC). The residual IL-23–dependent IL-17 immunity in homozygotes is sufficient to ensure antifungal immunity.

Like the *TYK2* P1104A variant, the *IL23R* R381Q variant is most frequent in populations of European ancestry (MAF: 0.06) (Boisson-Dupuis et al., 2018; Kerner et al., 2019; Kerner et al., 2021; Kerner et al., 2023). Moreover, these two alleles have evolved under negative selection in European populations over the last 4,500 years. Indeed, the frequency of *IL23R* R381Q has fallen from 10% in the Bronze age and 15% in the Neolithic to 5.5% today (Kerner et al., 2023). The frequency of *TYK2* P1104A has fallen from 10% in the Bronze age and 15% in the Neolithic to 2.9% today (Kerner et al., 2021; Kerner et al., 2023). These evolutionary trajectories are probably associated with the high burden of TB in Europeans between the Bronze age and the middle of the 20^th^ century (Kerner et al., 2021; Kerner et al., 2023). Both the *TYK2* P1104A and *IL23R* R381Q variants have a protective effect against inflammatory bowel diseases (ORs ranging from 0.1 to 0.3 and 0.2 to 0.7, respectively) (Boisson-Dupuis et al., 2018; Duerr et al., 2006; Momozawa et al., 2011). These observations suggest that the higher risk of inflammatory disorders in post-Neolithic Europeans may be due, in part, to the negative selection of alleles such as *TYK2* P1104A and *IL23R* R381Q that weaken protective immunity to primary infection by attenuating the inflammatory response (Boisson-Dupuis et al., 2018; Fieschi et al., 2003). The risk of TB is probably largely underestimated in our study, as few present-day Europeans are exposed to *M. tuberculosis*. The current clinical penetrance of partial IL-23R deficiency for TB in Europe is therefore probably low, and the impact of *IL23R* R381Q homozygosity is probably lower than that of *TYK2* P1104A at cellular level (Fig. 3 D). Genetic testing for these *IL23R* variants should nevertheless be offered to TB patients, especially those of European ancestry, as treatment with recombinant IFN-γ may be of therapeutic benefit (Casanova et al., 2024).

## Materials and methods

### Patients

Informed consent for participation in this study was obtained in accordance with local regulations, with approval from the institutional review board. The study was approved by the institutional ethics committees of The Rockefeller University and Necker Hospital for Sick Children and was performed in accordance with the requirements of these bodies. The experiments described here were performed in France and the United States of America, in accordance with local regulations.

### Whole-exome sequencing (WES), variant filtering, and Sanger sequencing

Genomic DNA was extracted from whole-blood samples from the patients and their relatives. WES was performed as previously described (Ogishi et al., 2022; Ogishi et al., 2025). MAFs in the general population, as reported in gnomAD database v4.1, and precomputed combined annotation–dependent depletion scores (v1.7) were used for variant filtering. The mutation significance cutoff was calculated as previously described (Itan et al., 2016). For the verification and familial segregation of variants, exons and flanking regions were amplified from DNA with DreamTaq DNA polymerase. They were then sequenced by the Sanger method with the Big Dye Terminator v3.1 kit (Thermo Fisher Scientific) and subjected to capillary electrophoresis (#A30469, Applied Biosystems 3500xL system, Thermo Fisher Scientific).

### Variant enrichment analysis

We performed an enrichment analysis on the G300V, R381Q, and G149R variants of IL-23R in our cohort of 1,874 TB patients and 20,883 healthy controls or patients with non-intramacrophagic infectious diseases. Association analyses were performed with regenie assuming a recessive mode of inheritance (Mbatchou et al., 2021). This method builds a whole-genome regression model based on common variants to correct for the effects of relatedness and population stratification. We used the approximate Firth P value when the logistic regression P value was below 0.05. A weight was applied to variants based on the ratio of WT to mutant protein activity to ensure that variants with more severe effects were given higher weights, as suggested by Wu et al. (2011). Analyses were adjusted for sex and the first 10 principal components (PC) of the PC analysis (PCA). PCA was performed with Plink v1.9 (Purcell et al., 2007) on WES data with a pruned set of 17,934 exonic variants (maximum linkage disequilibrium between each pair of SNPs <0.4) with a MAF >1% and a call rate >99%. We inferred genetic ancestry from the PCA, using a reference set of individuals, as previously described (Belkadi et al., 2016). Individuals with a prediction confidence >80% for similarity to individuals with Middle Eastern ancestry were retained for Middle East-specific analysis.

### Detection of founder effect

The age of the most recent common ancestor carrying the G300V variant was estimated using the EstiAge method (Genin et al., 2004), assuming a generation time of 27 years (Wang et al., 2023). In brief, the EstiAge method assumes that all affected individuals inherited the variant from a single common ancestor who introduced it generations ago. The number of generations is estimated from the length of the haplotype shared around the variant by locating the most likely recombination points on the ancestral haplotype across the different samples.

### Mammalian expression constructs

The generation of N-terminal NanoLuciferase-tagged WT IL-23R (NLuc IL-23R), HaloTag WT IL-12Rβ1 receptor (HaloTag IL-12Rβ1), and untagged IL-12Rβ1 constructs has been described elsewhere (Lay et al., 2022). The R381Q *IL23R* mutant was generated from the NL-IL23R plasmid with a Phusion site-directed mutagenesis kit (Thermo Fisher Scientific) according to the manufacturers’ instructions. The oligonucleotide primers used (Merck) were the following:

Forward primer 5′-ATTTAACAGATCATTCCAAACTGGGATTAAAAGAAGG-3′

Reverse primer 5′-ATCCCAATCAAAGAAAGAATTGACAAC-3′.

The PCR products were then digested with Dpn1 for 15 min at 37°C to eliminate methylated (template) DNA. The mutated R381Q *IL23* sequence was then excised from the PCR-generated plasmid vector and cloned back into the original template vector. Sequences were confirmed by Sanger sequencing (DEEPSEQ, University of Nottingham).

The NanoLuciferase-tagged *IL23R* plasmids containing mutations G300V and L372F were generated by site-directed mutagenesis with the Q5 Site-Directed Mutagenesis Kit (#E0554S; NEB) and the following primers:

G300V:

Forward primer: 5′-TCAAGAAACAGTGAAAAGGTACTGG-3′

Reverse primer: 5′-CATCTCACTTGAAATACGTAC-3′.

L372F:

Forward primer: 5′-AATTCTTTCTTTTATTGGGATATTTAACAG-3′

Reverse primer: 5′-GACAACATAACAGCAAAG-3′.

In all the above primer sequences, the underlined nucleotides were mutated.

The NanoLuciferase-tagged *IL23R* plasmid containing the G149R mutation was generated by site-directed mutagenesis and assembly cloning. Briefly, two separate PCR were performed with the primers below, with the WT NLuc *IL23* plasmid as a template. The PCR products were purified and assembled with the NEBuilder HiFi DNA Assembly Cloning Kit (#E5520S; NEB) according to the manufacturer’s instructions.

G149R:

Forward primer 1: 5′-CTGGAATGCTCGCAAGCTCACCTAC-3′

Reverse primer 1: 5′-GGAGCGAACGACCTACACCGAACTGAGATACCTACAGCG-3′

Forward primer 2: 5′-CGCTGTAGGTATCTCAGTTCGGTGTAGGTCGTTCGCTCC-3′

Reverse primer 2: 5′-GTAGGTGAGCTTGCGAGCATTCCAG-3′.

In all the above primer sequences, the underlined nucleotides were mutated.

All the NanoLuciferase-tagged *IL23R* receptor plasmids and variants of them were verified by whole-plasmid sequencing (Plasmidsaurus, London, using Oxford Nanopore Technology).

### Cell culture

PBMCs were isolated by Ficoll-Hypaque density centrifugation (Amersham Pharmacia Biotech). Human embryonic kidney cells (HEK293T [CRL3216; ATCC]) and EBV-B cells were cultured in DMEM or RPMI-1640 medium (Gibco), respectively, supplemented with 10% FCS (Sigma-Aldrich). For T-blast induction, PBMCs were cultured in ImmunoCult-XF T Cell Expansion Medium (STEMCELL) in the presence of ImmunoCultTM Human CD3/CD28/CD2 T cell activator (12.5 μl.ml^−1^) (STEMCELL) and human recombinant IL-2 (100 ng/ml, Novartis).

### RNA analysis and RT-qPCR

Total RNA was extracted with the Quick-RNA Microprep kit (Zymo) and reverse-transcribed to generate cDNA with the High-Capacity RNA-to-cDNA kit (Applied Biosystems). Quantitative PCR was then performed on the RNA with the Applied Biosystems probes/primers specific to IL23R-FAM (Hs00332759_m1) and β-glucuronidase-VIC (4326320E) for normalization. Results are expressed according to the ΔCt method.

### NanoBRET ligand-binding assays

HEK cells were used to seed a six-well plate (10578911; Corning Costar) at a density of 90,000 cells per well. The cells were incubated overnight at 37°C under an atmosphere containing 5% CO_2_. The next day, cells were transfected with a 4:1 ratio of WT or mutant NLuc IL-23R and IL-12Rβ1 cDNA with FuGENE HD (Promega Corporation), used according to the manufacturer’s instructions (3:1 total cDNA-to-reagent ratio). Cells were incubated overnight at 37°C under an atmosphere containing 5% CO_2_. The transfected cells were then plated at a density of 20,000 or 30,000 cells per well on poly-D-lysine–coated white, flat-bottomed 96-well plates (655098; Greiner Bio-One) and incubated overnight at 37°C under an atmosphere containing 5% CO_2_. The plating medium was then removed, and the cells were incubated with various concentrations of IL-23–TAMRA (Lay et al., 2022) in HEPES-buffered saline solution (HBSS; 2 mM sodium pyruvate, 145 mM NaCl, 10 mM D-glucose, 5 mM KCl, 1 mM MgSO_4_·7H_2_O, 10 mM HEPES, 1.3 mM CaCl_2_, and 1.5 mM NaHCO_3_ in double-distilled water, pH 7.45) supplemented with 0.1% BSA in the presence or absence of unlabeled IL-23 (Lay et al., 2022) (100 nM). IL-23–TAMRA was generated as previously described (Lay et al., 2022). Cells were incubated for 60 min at 37°C without CO_2,_ and the NanoLuciferase substrate furimazine was then added to all wells (Promega Corporation; final concentration of 7.7 μM). Plates were incubated for 5 min at room temperature, and the emitted luminescence and fluorescence were then simultaneously detected with a PHERAstar FS plate reader (BMG LabTech) fitted with a 450 ± 30 nm band-pass (luminescence emission [NLuc]) and >550 nm long-pass (fluorescence emission [IL-23–TAMRA]) filters.

### NanoLuciferase assays to quantify the cell surface expression of NLuc IL-23R

HEK cells were used to seed a 6-well plate (10578911; Corning Costar) at a density of 90,000 cells per well. They were incubated overnight at 37°C under an atmosphere containing 5% CO_2_. The cells were then transfected with a 4:1 ratio of WT or mutant NLuc *IL23R* and *IL12RB1* cDNA with FuGENE HD (Promega Corporation), according to the manufacturer’s instructions (3:1 cDNA-to-reagent ratio). Cells were incubated overnight at 37°C under an atmosphere containing 5% CO_2_ and were then used to seed poly-D-lysine–coated white, flat-bottomed 96-well plates (655098; Greiner Bio-One) at a density of 20,000 or 30,000 cells per well. The plates were incubated overnight at 37°C under an atmosphere containing 5% CO_2_. The culture medium was then removed and replaced with 7.7 μM of furimazine in HBSS (10 min at 37°C) in the presence or absence of NanoLuc extracellular inhibitor (60 μM; Promega Corporation). Luminescence emissions were then detected with a PHERAstar FS.

### Luminescence imaging

HEK cells were used to seed poly-D-lysine–coated 5-mm Cellview 4-quadrant culture dishes (Greiner Bio-one) with a 10-mm glass coverslip bottom at a density of 80,000 cells per quadrant. The plates were incubated overnight at 37°C under an atmosphere containing 5% CO_2_. The cells were then transfected with a 4:1 ratio of WT or mutant NLuc IL23R and HaloTag IL12RB1 cDNA with FuGENE HD (Promega Corporation), according to the manufacturer’s instructions (3:1 total cDNA-to-reagent ratio). Cells were incubated overnight at 37°C under an atmosphere containing 5% CO_2_. The plating medium was then removed and replaced with HBSS containing furimazine (1:400 final dilution). Luminescence images were acquired with an Olympus LuminoView 200 microscope fitted with a 60× NA1.42 oil immersion objective using a 0.5× tube lens and a C9100-23B IMAGE EMX2 camera (Hamamatsu, Japan) with an exposure time of 20 s (25 gain).

### NanoBRET intrareceptor interaction assays

HEK cells were used to seed a 6-well plate (10578911; Corning Costar) at a density of 90,000 cells per well. They were incubated overnight at 37°C under an atmosphere containing 5% CO_2_. The cells were then transfected with a 4:1 ratio of WT or mutant NLuc IL-23R and HaloTag IL-12Rβ1 cDNA with FuGENE HD (Promega Corporation), according to the manufacturer’s instructions (3:1 cDNA-to-reagent ratio), and incubated overnight at 37°C under an atmosphere containing 5% CO_2_. The transfected cells were used to seed poly-D-lysine–coated white, flat-bottomed 96-well plates (655098; Greiner Bio-One) at a density of 20,000–30,000 cell per well. They were incubated overnight at 37°C under an atmosphere containing 5% CO_2_. The culture medium was then removed and replaced with DMEM/10% FBS supplemented with the membrane-impermeant HaloTag 618 ligand (Promega Corporation) at a concentration of 100 nM. The cells were incubated for 5 h at 37°C under an atmosphere containing 5% CO_2_ and were then washed three times HBSS/0.1% BSA and incubated with HBSS/0.1% BSA containing 5 nM IL-23 for 60 min at 37°C. Furimazine was then added to all wells (7.7 μM final concentration), and the plate was allowed to stand for 5 min before the simultaneous measurement of luminescence and fluorescence emissions with a PHERAstar FS fitted with 460 ± 80 nm band-pass (luminescence [NLuc]) and >610 nm long-pass filters (fluorescence [HaloTag 618]).

### Mutagenesis for *IL23R* variants

Plasmids containing the WT human *IL23R* (#RG211477; OriGene) ORF were obtained, the tag was removed, and the variants studied here were generated by site-directed mutagenesis with specific primers and CloneAmp HiFi PCR premix (Takara). These ORFs included the *IL23R* S221F variant. The ORFs were introduced into the pLenti III-UbC-mCherry plasmid.

### Luciferase reporter assay

HEK cells were grown in DMEM supplemented with 10% FCS for 24 h before transfection. We assessed the impact of the mutation on IL-23R function by transfecting cells with the pLenti III-UbC-mCherry (empty vector or plasmid, EV) or the same plasmid backbone containing one of the *IL23R* ORFs (25 ng/well for a 96-well plate), pGL4.47(luc2P/SIE/Hygro) (Promega Corporation) reporter plasmids (100 ng/well), the pRL-SV40 plasmid (10 ng/well), and the PGK-*IL12RB1* plasmid (encoding WT IL-12Rβ1—50 ng/well) in the presence of X-tremeGENE 9 DNA Transfection Reagent (Roche). The medium was removed 36 h after transfection and replaced with DMEM supplemented with 10% FCS or the indicated cytokines for 16 h. Experiments were performed with technical duplicates, and the promoter activity of each well is expressed as firefly luciferase activity/*Renilla* luciferase activity normalized against the WT signal after stimulation with 10 ng/ml IL-23 taken as 100% activity.

### Assessment of STAT3 phosphorylation by western blotting

Levels of pSTAT3 were assessed in HEK or EBV-B cells. After 36 h of transfection (6 h with transfection medium followed by 30 h in DMEM containing 10% FCS), cells (1 × 10^6^) were incubated for 30 min with IL-23 (1 or 10 ng/ml, R&D) or IFN-α2a (1 ng/ml, Miltenyi Biotec). Cells were washed in PBS and lysed in RIPA lysis buffer (Thermo Fisher Scientific) supplemented with protease inhibitors (Roche mini EDTA-free, one tablet in 10 ml), phosphatase inhibitors (Roche PhosSTOP, one tablet in 10 ml), and benzonase (50 U/ml). Lysates were clarified and protein concentration was determined in a Bradford assay. Equal amounts of protein were subjected to SDS-PAGE, and the resulting bands were transferred to a PVDF membrane with 0.2 μm pores. Membranes were subjected to Ponceau staining, incubated in 5% skim milk in PBST for 1 h, briefly rinsed with PBST, and then incubated overnight at 4°C in primary antibody solution (5% BSA PBST or 5% skim milk PBST). Membranes were then washed three times, for 15 min each, in PBST, incubated in secondary antibody solution (1:2,000 dilution in 5% skim milk PBST) for 1 h at room temperature, and then washed again three times, for 15 min each, in PBST. Finally, chemiluminescence was detected with ECL reagents and a Bio-Rad ChemiDoc. After acquisition, the membrane was incubated in stripping buffer for 15 min (Restore Western Blot Stripping Buffer, Thermo Fisher Scientific) and rinsed in PBST. The induction of pSTAT3 induction is expressed as a percentage of the pSTAT3/STAT3 for the WT after stimulation with IL-23.

### Determination of IL-12Rβ1^+^ and mCherry^+^ cell frequency by flow cytometry

Cells (1 × 10^6^) were stained for 5 min at room temperature with the Aqua Dead cell viability marker (Thermo Fisher Scientific) and were then incubated for 30 min at 4°C with PE-conjugated anti-CD212 (IL-12Rβ1) antibody (Cat: 556065; BD Biosciences, Clone: 2.4E6, 1:5 dilution). The cells were analyzed in PE-Texas Red-, BV421-, and PE-compensated channels on an LSRFortessa X-20 flow cytometer (Beckman Coulter), and the results were analyzed with FlowJo software.

### Fresh PBMC stimulation

We used 100,000 PBMCs to seed RPMI supplemented with 10% human serum in each well of a round-bottomed 96-well plate. The cells were stimulated by incubation for 48 h with human recombinant IL-1β (2.5 ng/ml, R&D) and human recombinant IL-23 (0.001–100 ng/ml, R&D). PMA (Cat: P1585-1MG; Sigma-Aldrich, 8 ng/ml) + ionomycin (Cat: 56092-81-0; Sigma-Aldrich, 10-5 M) were added for the last 24 h only. The supernatants were collected and subjected to LEGENDplex multiplex ELISA with Human Inflammation Panel 1 (740809; BioLegend), according to the manufacturer’s instructions.

### PBMC stimulation with BCG

Freshly thawed PBMCs from healthy controls, *IL23R*^G300V/G300V^, *IL23R*^R381Q/R381Q^, *IL23R* ^G149R/G149R^, *TYK2*^P1104A/P1104A^, IL-12Rβ1-, and IL23R-deficient patients were dispensed into a U-bottomed 96-well plate at a density of 3 × 10^5^ cells per well, in 200 μl lymphocyte medium per well, as previously described (Philippot et al., 2023; Yang et al., 2020). Cells were incubated in the presence or absence of live BCG, at a multiplicity of infection of 1, with or without recombinant human IL-12 (5 ng/ml, R&D Systems), recombinant human IL-18 (25 ng/ml, R&D Systems), and/or recombinant human IL-23 (100 ng/ml, 1290-IL R&D Systems) or IFN-γ (5,000 IU/ml). After 40 h of stimulation, GolgiPlug (555029; BD Biosciences; 1:1,000 dilution) was added to each well to inhibit cytokine secretion. Cells were also costimulated for 24 h with IL-23 (100, 10, or 1 ng/ml) and IL-18 (200 ng/ml). PMA ionomycin was added 6 h before the end of incubation. The cells were collected by centrifugation for flow cytometry analysis. They were stained with the Zombie NIR Fixable Viability Kit (BioLegend; 1:2,000 dilution) at room temperature for 15 min and then stained on ice for 30 min with a surface-staining panel containing FcR blocking reagent (Miltenyi Biotec; 1:50 dilution), anti-CD3-Alexa Fluor 532 (Clone: UCHT1, 58-0038-42; eBioscience; 1:50 dilution), anti-γδTCR-FITC (Clone:B1.1, 11-9959-41; eBioscience; 1:50 dilution), anti-Vδ2-APC/Fire 750 (Clone:B6, 331419; BioLegend; 1:100 dilution), anti-CD56-BV605 (Clone: 5.1H11, 362537; BioLegend; 1:100 dilution), anti-CD4-BV750 (Clone: SK3, 566356; BD Biosciences; 1:800 dilution), anti-CD8a-Pacific Blue (Clone: SK1, 344717; BioLegend; 1:100 dilution), anti-Vα7.2 TCR-APC (Clone: 3C10, 351708; BioLegend; 1:100 dilution), anti-Vα24-Jα18-PE/Cy7 Clone: 6B11, 342912; BioLegend; 1:100 dilution), anti-CD20-BV785 (Clone: 2H7, 302356; BioLegend; 1:200 dilution), and anti-PD-1-PE (Clone: MIH4, 12-9969-42; eBioscience; 1:100 dilution) antibodies. Cells were fixed by incubation with 2% paraformaldehyde in PBS on ice for 15 min. They were then permeabilized/stained by incubation overnight at −20°C in the permeabilization buffer from the Nuclear Transcription Factor Buffer Set (BioLegend) with an intracellular cytokine panel containing FcR blocking reagent (Miltenyi Biotec; 1:50 dilution), anti-IFN-γ-BV711 (Clone: 4 S.B3, 502540; BioLegend; 1:50 dilution), anti-TNF-BV510 (Clone: MAb11, 502950; BioLegend; 1:50 dilution), and anti-IL-10-PE/Dazzle594 (Clone: JES3-19F1, 506812; BioLegend; 1:50 dilution) antibodies. As a positive control, cells in a separate well were stimulated by incubation with PMA (Sigma-Aldrich; 25 ng/ml) and ionomycin (Sigma-Aldrich; 500 nM) for 1 h without GolgiPlug and then for 7 h with GolgiPlug (for intracellular cytokine staining). Cells were acquired with an Aurora cytometer (Cytek). Data were manually gated with FlowJo.

### Mass cytometry immunophenotyping

Deep immunophenotyping was performed by mass cytometry (CyTOF). We used 200 μl fresh whole blood from the patients and controls. We used a previously described custom-designed panel (Momenilandi et al., 2024), according to Standard BioTools’s instructions. Cells were frozen at[−80°C after overnight dead-cell staining, and acquisition was performed on a Helios mass cytometer (Fluidigm). All the samples were processed within 24–36 h of sampling. Data analysis was performed with OMIQ software.

### PBMC stimulation for scRNAseq

PBMCs were thawed and plated at a density of 50,000–300,000 cells per well in a U-bottom 96-well plate. IL-23 was added to a final concentration of 100 ng/ml. After 6 h of incubation at 37°C, non-stimulated and stimulated cells were recovered, washed three times in PBS supplemented with 0.5% FBS, filtered through a 40-µm-pore MACS strainer, and subjected to scRNAseq.

### Analysis of scRNAseq data

For baseline analysis, scRNAseq data were integrated with previously published data for eight controls and data for three newly studied healthy controls, together with samples from IL-12Rβ1^−/−^ (*n* = 3), TYK2^P1104/P1104^ (*n* = 3), IL-23R^−/−^ (*n* = 3), IL-23R ^R381Q/R381Q^ (*n* = 3), IL-23R^G300V/G300V^ (*n* = 2), and IL-23R^G149R/G149R^ (*n* = 1) patients. For the stimulation analysis, paired non-stimulated and IL-23–stimulated samples (data for the 6-h time point) were analyzed, including six previously published adult controls, two newly studied healthy controls, and patient samples: IL-12Rβ1^−/−^ (*n* = 1), TYK2^P1104/P1104^ (*n* = 2), IL-23R^−/−^ (*n* = 2), IL-23R^R381Q/R381Q^ (*n* = 3), IL-23R^G300V/G300V^ (*n* = 2), and IL-23R^G149R/G149R^ (*n* = 1).

Quality control was performed by filtering cells based on the percent mitochondrial gene expression, the total number of transcripts detected (UMIs), and the number of genes expressed per cell, with standard thresholds. Cell type annotation was performed by anchor-based label transfer in Seurat (Hao et al., 2021). A previously integrated PBMC reference (canonical correlation analysis, CCA) was processed by PCA and UMAP. Anchors were identified between the CCA-integrated reference and PCA-transformed query data, making it possible to transfer cell type labels and projections onto the reference UMAP space. UMAP was used for visualization. Gene expression was quantified in Seurat (Hao et al., 2021). Pseudobulk expression profiles were generated by summing counts per sample and cell type with the SingleCellExperiment framework (Amezquita et al., 2020) and muscat (Crowell et al., 2020). PCA was performed on variance-stabilized counts, with batch effects corrected with removeBatchEffect from limma (Ritchie et al., 2015). Differential expression analysis was performed with DESeq2 (Love et al., 2014), and log_2_ fold-change shrinkage for gene ranking was applied with apeglm (Zhu et al., 2019). GSEA was performed with fgsea (Korotkevich et al., 2021, *Preprint*) and gene sets from the Molecular Signatures Database (Liberzon et al., 2011).

We calculated the induction of *IFNG* mRNA following IL-23 stimulation using the housekeeping genes HKG, C1orf43, CHMP2A, EMC7, GPI, PSMB2, PSMB4, RAB7A, REEP5, SNRPD3, VCP, and VPS29, with geometric mean normalization for each sample.

### Statistical analysis

Statistical analyses were performed with GraphPad Prism v8.4.3 and v10.5.0. Paired *t* tests (intrareceptor BRET experiments), one-way ANOVA with Tukey’s post hoc correction for multiple comparisons (IL-23R–IL12Rβ1 interaction experiments), and nonparametric Mann–Whitney *U* tests were used as appropriate to assess statistical significance. A P value <0.05 was considered statistically significant, *; P < 0.01, **; P < 0.001, ***; and P < 0.0001, ****. PCA was performed with Plink v1.9 software on WES data, using 16,730 exonic variants with a MAF >1% and a call rate >98%.

BRET ratios were calculated by dividing fluorescence by luminescence emissions. Pharmacological data analysis was performed with GraphPad Prism 10.1.0. IL-23–TAMRA ligand-binding data were fitted by nonlinear regression with the equation

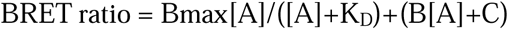

where “Bmax” is the maximum specific binding BRET signal, “[A]” is the concentration of IL-23–TAMRA, “B” is the slope of the nonspecific binding component, and “C” is the *y* intercept. The specific binding of IL-23–TAMRA was determined by subtracting the BRET ratios measured in the presence of unlabeled IL-23 from those measured in its absence. Specific binding data were fitted with the equation

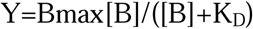

where Bmax is the maximum specific binding signal of the curve and [B] is the concentration of IL-23–TAMRA used. The K_D_ values presented are the mean ± SEM for each individual experiment.

For cell surface expression experiments, the emitted luminescence measured in the presence of the extracellular NanoLuciferase inhibitor is expressed as a percentage of that measured in the absence of the inhibitor (100%). For IL-12Rβ1 interaction experiments, WT NLuc IL-23R/HaloTag IL-12Rβ1 responses in the absence of IL-23 were set to 100%, and subsequent BRET ratios were normalized against this value and expressed as a percentage ± SEM.

### Online supplemental material

Fig. S1 shows IL12RB1 and IL23R expression at the mRNA and protein levels in transfected HEK cells, along with STAT3 phosphorylation after IL-23 stimulation in cells expressing different IL23R variants. Fig. S2 displays the predicted population-level impact of IL23R variants based on AlphaMissense, as well as their functional effects on IL-23 binding and STAT3 phosphorylation assessed by western blotting in homozygous EBV-B cells. These EBV-B cell lines were generated from the individuals mentioned. PBMC immunophenotyping of individuals homozygous for hypomorphic IL23R variants is shown in Fig. S3. Analysis of the single-cell sequencing of PBMCs from individuals homozygous for hypomorphic IL23R variants is shown in Fig. S4. Fig. S5 shows impaired IFN-γ induction in PBMCs from individuals homozygous for hypomorphic IL23R variants. Table S1 reports *IL23R* variants in the diverse cohorts of patients, and Table S2 shows the association in the HGID cohort for G149R, G300V, and R381Q variants (no homozygotes for L372F were found in the HGID cohort) for the recessive model.

## Acknowledgments

We thank the patients and their families for placing their trust in us. We thank the members of both branches of the Laboratory of Human Genetics of Infectious Diseases. We thank Tatiana Kochetkov for technical assistance; Yelena Nemirovskaya, Mark Woollett, Dana Liu, Amyrat Geraldo, Soraya Boucherit, Abigail Bejou, Maya Chrabieh, and Lazaro Lorenzo for administrative assistance. We thank the Vecteurs Viraux et Transfert de Gènes (VVTG) platform of the “Necker Enfants Malades” Institute for the EBV-B cell line. We thank the Flow Cytometry Resource Center at The Rockefeller University. We thank the National Institutes of Health (NIH) Tetramer Core Facility for providing the MR1 tetramer, which was developed jointly with Dr. James McCluskey, Dr. Jamie Rossjohn, and Dr. David Fairlie. We are indebted to the “Biobanc de l’Hospital Infantil Sant Joan de Déu per a la Investigación” a member of the ISCIII National Network of Biobanks, Spain, for sample and data procurement. Graphical abstract was created with Biorender.com (https://app.biorender.com/citation/69d3f0d806893ae4a332727b).

The HGID laboratory was funded in part by the National Institute of Allergy and Infectious Diseases, NIH (grant numbers R01AI095983 to J.-L. Casanova and J. Bustamante and R01AI127564 to J.-L. Casanova and A. Puel), the National Center for Research Resources and the National Center for Advancing Sciences, NIH (grant number UL1TR001866 for J.-L. Casanova), The Rockefeller University, the St. Giles Foundation, the Howard Hughes Medical Institute, Institut National de la Santé et de la Recherche Médicale (INSERM), Paris Cité University, the French Agence Nationale de la Recherche (ANR) under the France 2030 program (ANR-10-IAHU-01), the Integrative Biology of Emerging Infectious Diseases Laboratory of Excellence (ANR-10-LABX-62-IBEID), the ANR project MAFMACRO (ANR-22-CE92-0008 to J. Bustamante), the Agence Nationale de Recherches sur le Sida et les Hépatites Virales (ANRS) projects ECTZ170784-ANRS0073 and ANRS0787-ECTZ323676 to S. Boisson-Dupuis, the French Foundation for Medical Research (FRM) (EQU202503020018), the European Union’s Horizon 2020 Research and Innovation Program under grant agreement no. 824110 (EASI-genomics), the Square Foundation, Grandir -Fonds de Solidarité pour L’enfance, William E. Ford, General Atlantic’s Chairman and Chief Executive Officer, Gabriel Caillaux, General Atlantic’s Co-President, Managing Director, and Head of Business at EMEA, and the General Atlantic Foundation. C. Conil was supported by an ANRS PhD fellowship (ECTZ293125), and Q. Philippot was supported by the Assistance Publique Hôpitaux de Paris (Année Recherche) and by the MD-PhD program of INSERM (Ecole de l’INSERM Liliane Bettencourt). J. Bohlenwas supported by fellowships from the European Molecular Biology Organization and Marie Curie Research. E. Feredj was supported by the FRM PhD program, Poste de Thèse Pour Interne et Assistant (FDM202406018939), “bourse d’aide à la mobilité” de l’Institut Servier, Collège des Universitaires des Maladies Infectieuses et Tropicales, European Society of Clinical Microbiology and Infectious Diseases, and “bourse de mobilité internationale” Réunion Interdisciplinaire de Chimiothérapie Anti-Infectieuse. H. Li was supported by Labex-IBEID and an ANRS PhD fellowship (ECTZ293911). M. Momenilandi was supported by an Imagine Institute Postdoctoral Prize. M. Ogish was supported by the David Rockefeller Graduate Program, the Funai Foundation for Information Technology, the Honjo International Scholarship Foundation, and the New York Hideyo Noguchi Memorial Society. S.J. Hill, L.E. Kilpatrick, and S. Platt are funded by the Medical Research Council, UK (grant no. MR/W016176/1). C.S. Lay was supported by a Biotechnology and Biological Sciences Research Council Industrial Case Studentship with GSK to C.S. Lay, P.D. Craggs, and S.J. Hill (grant no. BB/S507027/1). This study was supported by the project PI15/01094 to L. Abel, integrated in the Plan Nacional de I+D+I and cofinanced by the ISCIII – Subdirección General de Evaluación y Formento de la Investigación Sanitaria – and the Fondo Europeo de Desarrollo Regional. I. Meyts is funded by Fonds Wetenschappelijk Onderzoek (FWO) Vlaanderen G1805826N. A.A. Arias, M. Tejada - Giraldo, A. Baena, and L.F. Barrera were supported by the Ministerio de Ciencia, Tecnología e Innovación MINCIENCIAS (111584467551/CT 415-2020), the Jeffrey Modell Foundation, and the Fundación Diana García de Olarte para las Inmunodeficiencias Primarias (Colombia). C.S. Ma and S.G. Tangye are supported by Investigator Grants awarded by the National Health and Medical Research Council of Australia (C.S. Ma: 2017463 [level 1]; S.G. Tangye: 1176665 [level 3]).

## Author contributions

Diana Olguín Calderón: conceptualization, data curation, formal analysis, investigation, methodology, validation, visualization, and writing—original draft, review, and editing. Laura E. Kilpatrick: formal analysis, funding acquisition, investigation, visualization, and writing—original draft, review, and editing. Clément Conil: data curation, formal analysis, software, validation, and writing—original draft, review, and editing. Quentin Philippot: conceptualization, formal analysis, investigation, and writing—original draft, review, and editing. Masato Ogishi: investigation, methodology, software, and writing—review and editing. Joseph Vellutini: data curation, formal analysis, resources, software, and visualization. Ji Eun Han: methodology, resources, and validation. Narelle Keating: investigation and visualization. Hailun Li: investigation. Geetha Rao: formal analysis and investigation. Jonathan Bohlen: conceptualization, investigation, methodology, supervision, validation, and writing—review and editing. Charles S. Lay: investigation and methodology. Simon Platt: investigation, methodology, and writing—original draft. Gaspard Kerner: formal analysis. Elsa Feredj: investigation. Jessica N. Peel: writing—review and editing. Mana Momenilandi: investigation and resources. Yoann Seeleuthner: data curation and resources. Candice Lainé: investigation and resources. Camille Soudée: investigation. Claire Leloup: investigation. Cecile Debuisson: resources. Fanny Lanternier: investigation and visualization. Samuel Bitoun: data curation, formal analysis, investigation, validation, and writing—review and editing. Stephan Pavy: resources. Xavier Mariette: resources and writing—review and editing. Aniss Rafik: conceptualization, investigation, and resources. Hanaa Skhoun: conceptualization, investigation, and resources. Hanane El Ouazzani: conceptualization, investigation, and resources. Ismael Abderahmani-Ghorfi: conceptualization, investigation, and resources. Jamila El-Baghdadi: conceptualization, investigation, and resources. Andrés Baena: resources. Manuela Tejada -Giraldo: investigation. Luis Fernando Barrera: investigation and resources. Andrés Augusto Arias: investigation and resources. Giovanna Fabio: resources and visualization. Maria Carrabba: investigation and resources. Melike Emiroglu: data curation, resources, and supervision. Liliana Bezrodnik: resources. Loubna El Zein: investigation and resources. Hassan Hammoud: resources. Peter K. Gregersen: data curation. Benjamin Terrier: data curation and writing—review and editing. Rafael Leon Lopez: resources. Marion Touzet: investigation. Vincent Pestre: investigation and resources. Marlène Pasquet: resources. Lars Rogge: investigation, resources, and writing—review and editing. Michael Fayon: investigation and validation. François Galode: resources. Eric Jeziorski: resources. Darragh Duffy: resources and writing—review and editing. Lluis Quintana-Murci: resources and validation. Etienne Patin: resources, Charlotte Cunningham-Rundles: resources. Isabelle Meyts: data curation, funding acquisition, investigation, resources, and writing—review and editing. Shen-Ying Zhang: resources and writing—review and editing. Qian Zhang: resources. Emmanuelle Jouanguy: resources and writing—review and editing. Bertrand Boisson: resources. Jérémie Rosain: resources. Vivien Béziat: formal analysis. Mohammad Shahrooei: conceptualization, data curation, formal analysis, investigation, and validation. Seyed Alireza Mahdaviani: conceptualization, investigation, project administration, validation, and writing—review and editing. Nima Rezaei: conceptualization, data curation, investigation, methodology, project administration, supervision, validation, and writing—review and editing. Nima Parvaneh: investigation and resources. Zahra Chavoshzadeh: resources. Niloufar Yazdanpanah: resources. Nathalie Aladjidi: resources. Antoni Noguera-Julian: investigation and writing—review and editing. Ana Esteve-Solé: investigation, resources, and writing—review and editing. Laia Alsina: data curation, investigation, validation, and writing—review and editing. Davood Mansouri: resources. Sevgi Keles: data curation and investigation. Mediha Gonenc Ortakoylu: Conceptualization, data curation, formal analysis, funding acquisition, investigation, methodology, project administration, resources, software, supervision, validation, visualization, and writing—original draft, review, and editing. Deniz Aygun: data curation. Esra Yucel: resources. Ayca Kiykim: resources and writing—review and editing. Yildiz Camcioglu: data curation. Cindy S. Ma: funding acquisition, methodology, supervision, and writing—review and editing. Stuart G. Tangye: funding acquisition, project administration, supervision, validation, and writing—review and editing. Peng Zhang: data curation, formal analysis, and software. Laurent Abel: formal analysis and writing—review and editing, Peter D. Craggs: funding acquisition and project administration. Jean-Laurent Casanova: conceptualization, funding acquisition, project administration, resources, supervision, validation, and writing—original draft, review, and editing. Aurélie Cobat: methodology, supervision, validation, and writing—review and editing. Anne Puel: conceptualization, funding acquisition, resources, supervision, and writing—original draft, review, and editing. Jacinta Bustamante: conceptualization, formal analysis, funding acquisition, investigation, methodology, resources, supervision, and writing—original draft, review, and editing. Stephen J. Hill: funding acquisition, investigation, visualization, and writing—original draft, review, and editing. Stéphanie Boisson-Dupuis: conceptualization, data curation, formal analysis, funding acquisition, investigation, project administration, resources, supervision, validation, visualization, and writing—original draft, review, and editing.

## CRediT Role(s)

**DianaOlguín Calderón:**ConceptualizationData curationFormal analysisInvestigationMethodologyValidationVisualizationWriting - original draftWriting - review & editing

**Laura E.Kilpatrick:**Formal analysisFunding acquisitionInvestigationVisualizationWriting - original draftWriting - review & editing

**ClémentConil:**Data curationFormal analysisSoftwareValidationWriting - original draftWriting - review & editing

**QuentinPhilippot:**ConceptualizationFormal analysisInvestigationWriting - original draftWriting - review & editing

**MasatoOgishi:**InvestigationMethodologySoftwareWriting - review & editing

**JosephVellutini:**Data curationFormal analysisResourcesSoftwareVisualization

**Ji EunHan:**MethodologyResourcesValidation

**NarelleKeating:**InvestigationVisualization

**HailunLi:**Investigation

**GeethaRao:**Formal analysisInvestigation

**JonathanBohlen:**ConceptualizationInvestigationMethodologySupervisionValidationW riting - review & editing

**Charles S.Lay:**InvestigationMethodology

**SimonPlatt:**InvestigationMethodologyWriting - original draft

**GaspardKerner:**Formal analysis

**ElsaFeredj:**Investigation

**Jessica N.Peel:**Writing - review & editing

**ManaMomenilandi:**InvestigationResources

**YoannSeeleuthner:**Data curationResources

**CandiceLainé:**InvestigationResources

**CamilleSoudée:**Investigation

**ClaireLeloup:**Investigation

**CecileDebuisson:**Resources

**FannyLanternier:**InvestigationVisualization

**SamuelBitoun:**Data curationFormal analysisInvestigationValidationWriting - review & editing

**StephanPavy:**Resources

**XavierMariette:**ResourcesWriting - review & editing

**AnissRafik:**ConceptualizationInvestigationResources

**HanaaSkhoun:**ConceptualizationInvestigationResources

**HananeEl Ouazzani:**ConceptualizationInvestigationResources

**IsmaelAbderahmani-Ghorfi:**ConceptualizationInvestigationResources

**JamilaEl-Baghdadi:**ConceptualizationInvestigationResources

**AndrésBaena:**Resources

**ManuelaTejada-Giraldo:**Investigation

**Luis FernandoBarrera:**InvestigationResources

**Andrés AugustoArias:**InvestigationResources

**GiovannaFabio:**ResourcesVisualization

**MariaCarrabba:**InvestigationResources

**MelikeEmiroglu:**Data curationResourcesSupervision

**LilianaBezrodnik:**Resources

**LoubnaEl Zein:**InvestigationResources

**HassanHammoud:**Resources

**Peter K.Gregersen:**Data curation

**BenjaminTerrier:**Data curationWriting - review & editing

**RafaelLeon Lopez:**Resources

**MarionTouzet:**Investigation

**VincentPestre:**InvestigationResources

**MarlènePasquet:**Resources

**LarsRogge:**InvestigationResourcesWriting - review & editing

**MichaelFayon:**InvestigationValidation

**FrançoisGalode:**Resources

**EricJeziorski:**Resources

**DarraghDuffy:**ResourcesWriting - review & editing

**LluisQuintana-Murci:**ResourcesValidation

**EtiennePatin:**Resources

**CharlotteCunningham-Rundles:**Resources

**IsabelleMeyts:**Data curationFunding acquisitionInvestigationResourcesWriting - review & editing

**Shen-YingZhang:**ResourcesWriting - review & editing

**QianZhang:**Resources

**EmmanuelleJouanguy:**ResourcesWriting - review & editing

**BertrandBoisson:**Resources

**JérémieRosain:**Resources

**VivienBéziat:**Formal analysis

**MohammadShahrooei:**ConceptualizationData curationFormal analysisInvestigationValidation

**Seyed AlirezaMahdaviani:**ConceptualizationInvestigationProject administrationValidationWriting - review & editing

**NimaRezaei:**ConceptualizationData curationInvestigationMethodologyProject administrationSupervisionValidationWriting - review & editing

**NimaParvaneh:**InvestigationResources

**ZahraChavoshzadeh:**Resources

**NiloufarYazdanpanah:**Resources

**NathalieAladjidi:**Resources

**AntoniNoguera-Julian:**InvestigationWriting - review & editing

**AnaEsteve-Solé:**InvestigationResourcesWriting - review & editing

**LaiaAlsina :**Data curationInvestigationValidationWriting - review & editing

**DavoodMansouri:**Resources

**SevgiKeles:**Data curationInvestigation

**Mediha GonencOrtakoylu:**ConceptualizationData curationFormal analysisFunding acquisitionInvestigationMethodologyProject administrationResourcesSoftwareSupervisionValidationVisualizationWriting - original draftWriting - review & editing

**DenizAygun:**Data curation

**EsraYucel:**Resources

**AycaKiykim:**ResourcesWriting - review & editing

**YildizCamcioglu:**Data curation

**Cindy S.Ma:**Funding acquisitionMethodologySupervisionWriting - review & editing

**Stuart G.Tangye:**Funding acquisitionProject administrationSupervisionValidationWriting - review & editing

**PengZhang:**Data curationFormal analysisSoftware

**LaurentAbel:**Formal analysisWriting - review & editing

**Peter D.Craggs:**Funding acquisitionProject administration

**Jean-LaurentCasanova:**ConceptualizationFunding acquisitionProject administrationResourcesSupervisionValidationWriting - original draftWriting - review & editing

**AurélieCobat:**MethodologySupervisionValidationWriting - review & editing

**AnnePuel:**ConceptualizationFunding acquisitionResourcesSupervisionWriting - original draftWriting - review & editing

**JacintaBustamante:**ConceptualizationFormal analysisFunding acquisitionInvestigationMethodologyResourcesSupervisionWriting - original draftWriting - review & editing

**Stephen J.Hill:**Funding acquisitionInvestigationVisualizationWriting - original draftWriting - review & editing

**StéphanieBoisson-Dupuis:**ConceptualizationData curationFormal analysisFunding acquisitionInvestigationProject administrationResourcesSupervisionValidationVisualizationWriting - original draftWriting - review & editing

## Data availability

The data underlying the figures are available in the published article and its online supplemental material. In addition, scRNAseq data are available with the following BioProject ID: <u>PRJNA1449438</u> (https://www.ncbi.nlm.nih.gov/bioproject/PRJNA1449438).

## Non-standard abbreviation

IL: interleukin
IFN-γ: Interferon gamma
NK: Natural Killer
MAF: Minor allele frequency
MSMD: Mendelian susceptibility to mycobacterial diseases
BCG: bacille Calmette et Guérin
MAIT: Mucosal-associated invariant T cells
TB: Tuberculosis
UbC: Ubiquitin C
CMV: Cytomegalovirus
HGID: Human Genetics of Infectious Diseases
EBV-B: Epstein Barr virus-immortalized B cells
CMC: Chronic mucocutaneous candidiasis

## Legends for the supplementary figures

**Supplementary figure 1:** Analysis of *IL23R* variants from the general population. (A) Frequency of mCherry^+^ and of IL-12Rβ1^+^ cells in the mCherry^+^ population and levels of *IL23R* and *IL12RB1* mRNA in HEK293T cells transiently transduced with plasmids encoding IL-12Rβ1 and the various IL-23R variants expressed together with the mCherry reporter gene and a luciferase reporter. Order of plasmids is as follow: NT, EV, followed by IL23R WT, C115Y, G300V, R381Q, Q3H, A55T, R86Q, G149R, K150T, V159M, V160A, L193F, A199V, S221F, P306S, L30P, V362I, G300V+V362I, L372F, I373F, Q487H, and S559R. Bars indicate the mean ± SD for three independent biological replicates. (B) Detection by western blotting of pSTAT3 after IL-23 (10 ng/ml) and IFN-α (1 ng/ml) stimulation in HEK293T cells transfected with the indicated hypomorphic *IL23R* alleles, with total STAT3 and vinculin as loading controls. At least two independent experiments for each variant were performed. The blot is duplicated in Fig. 1 B. Source data are available for this figure: SourceData FS1.

**Supplementary figure 2:** Population genetics and impact of hypomorphic *IL23R* variants on IL-23 binding. **(A)** AlphaMissense/MAF (right) graphs for all homozygous *IL23R* LOF variants described in previous studies (red) (Martinez-Barricarte et al., 2018; Philippot et al., 2023), from the general population (gray dots), and other homozygous variants found in our in-house cohort (black triangles). The four hypomorphic variants are presented in different colors. MSC, mutation significance cutoff score with a 99% CI. CADD, combined annotation-dependent depletion. **(B)** Same graph as Fig. 2 C with an adjusted y axis, illustrating the saturable binding of IL-23–TAMRA to NLuc IL23R G300V/IL-12Rβ1, but with a low maximum BRET window. **(C)** HEK cells were transiently transfected with N-terminal NanoLuciferase-tagged versions of the WT or variants of the IL-23 receptor (NLuc IL-23R, NLuc IL-23R R381Q, NLuc IL-23R G300V, NLuc IL-23R G149R, or NLuc IL-23R L372F) and N-terminal HaloTag IL-12Rβ1. Emitted luminescence and fluorescence emissions were measured with a BMG PheraStar FS. The data shown are the mean ± SEM from five independent experiments performed in triplicate wells. Statistical significance was determined by one-way ANOVA with Tukey’s post hoc analysis for multiple comparisons with **P < 0.01, ***P < 0.001. **(D)** Western blot of pSTAT3 after the stimulation with 1 ng/ml or 10 ng/ml IL-23 stimulation of EBV–B cell from healthy controls (green), *IL23R*^R381Q/R381Q^ (violet), *IL23R*^G300V/G300V^ (orange), *IL23R*^G149^ ^R/G149R^, and *IL23R* LOF (red) homozygotes. IFN-α stimulation was used as a positive control. The representative of at least two independent experiment is showed. **(E)** Pedigrees of the kindreds studied in this report. The gene and variants are indicated above the kindreds. The clinical phenotype of each symptomatic patient is indicated by different shaded shapes, as described at the bottom of the figure. Symbols linked with a double line indicate consanguinity. The genotype is indicated under each symbol, with M corresponding to the variant found in each kindred. Arrows indicate the index case in each family. E?: Unavailable clinical or genotype information. Source data are available for this figure: SourceData FS2.

**Supplementary Figure 3:** Normal leukocyte development in patients homozygous for hypomorphic *IL23R* variants. **(A)** Frequencies and absolute counts of NK, γδ T, and MAIT cells, and frequency of T_H_1, T_H_2, T_H_17, and T_H_1* memory CD4^+^ T cells, in healthy adult (*n* = 35) and pediatric (*n* = 11) controls, *IL23R*^G149R/G149R^ (*n* = 1), *IL23R*^G300V/G300V^ (*n* = 1), *IL23R*^R381Q/R381Q^ (pediatric patients <18 years old [*n* = 6] and adults >18 years old [*n* = 3]), and *IL23R*^−/−^ (*n* = 2) individuals. **(B)** Frequencies of CD4^+^, CD8^+^, monocytes, and dendritic cells in healthy adults (*n* = 35) and pediatric (*n* = 11) controls, adult (*n* = 3), and pediatric (*n* = 6) homozygotes for R381Q, an adult homozygous for G149R, and an adult homozygous for G300V IL-23R-deficient patients, as assessed by spectral flow cytometry on fresh PBMCs. The statistical significance of differences was assessed in unpaired Mann–Whitney *U* tests with *P < 0.05, **P < 0.01. **(C)** Absolute counts of CD4^+^, CD8^+^, and NK subsets; monocytes, and dendritic cells (DCs) in healthy adults (*n* = 35) and pediatric (*n* = 11) controls, adult (*n* = 3) and pediatric (*n* = 6) patients homozygous for R381Q, one adult homozygous for G300V, and an adult homozygous for G149R IL-23R–deficient patients, assessed by CyTOF cytometry on fresh PBMCs. **(D)** Frequencies of T_H_1, T_H_2, T_H_17, and T_H_1* memory CD4^+^ T cells and absolute counts of NK, γδ T cells, CD4^+^, CD8^+^, and NK subset; monocytes, and dendritic cells (DCs) in healthy adults (*n* = 19) and pediatric (*n* = 20) controls, *IL23R*^G300V/G300V^ (*n* = 1), *IL23R*^R381Q/R381Q^ (pediatric patients <18 years old [*n* = 2] and adults >18 years old [*n* = 1]), *IL23R*^−/−^ (*n* = 3) and *IL12RB1*^−/−^ (*n* = 1) individuals, as determined with a spectral flow panel not recognizing MAIT cells.

**Supplementary figure 4:** Single-cell sequencing analysis of PBMCs from IL-23R-deficient patients. **(A)** UMAP representation of scRNAseq performed on PBMCs from 11 healthy controls, 3 *IL23R*^R381Q/R381Q^, 2 *IL23R*^G300V/G300V^, 1 *IL23R*^G149R/G149R^, 3 *IL23R*^−/−^, 3 *IL12R*β*1*^−/−^, and 3 *TYK2*^P1104A/P1104A^ patients. **(B)** Proportions of leukocyte subsets from the individuals in A at steady state. **(C)** Relative abundance of each leukocyte subset according to clustering analysis for non-stimulated and IL-23–stimulated samples. **(D)** Two-dimensional plots of IL-23–induced delta *IFNG* expression. The fold-change difference in *IFNG* mRNA levels following IL-23 stimulation in leukocytes from *IL23R*^R381Q/R381Q^, *IL23R*^G300V/G300V^, *IL23R*^G149R/G149R^, and *TYK2*^P1104A/P1104A^ patients relative to controls is shown on the x-axis. The y-axis shows the same parameter for IL-12Rβ1^−/−^ patients as a comparison. The color of the circles indicates the median change (IL-23 versus NS) in normalized *IFNG* mRNA levels in controls for the corresponding subsets. MAIT, NK, and Vδ2^+^ γδT cells are highlighted.

**Supplementary figure 5:** Impaired IFN-g induction in PBMCs from individuals homozygous for hypomorphic *IL23R* variants. **(A–C)** Freshly thawed PBMCs of the indicated genotypes were stimulated with IL-23 (100 ng/ml) in the presence or absence of IL-18 (25 ng/ml) for 48 h. Percentages of IFN-γ^+^ cells were assessed: (A) among MAIT cells, (B) among Vδ2^+^ γδ T cells, (C) among NK cells. Percentages of IFN-γ^+^ and TNF^+^ cells after PMA/ionomycin were evaluated as controls and are shown in Fig. S5, G–I. Healthy (*n* = 16) controls, *IL23R*^G149R/G149R^ (*n* = 1), *IL23R*^G300V/G300V^ (*n* = 5), *IL23R*^R381Q/R381Q^ (*n* = 6), *IL12R*β*1*^−/−^ (*n* = 1), and *TYK2*^P1104A/P1104A^ (*n* = 5) individuals were investigated. Nonparametric Mann–Whitney *U* tests were used for analysis, with *P < 0.05, **P < 0.01, ***P < 0.001, ****P < 0.0001. **(D)** Fresh PBMCs of the indicated genotypes were left unstimulated (NS) or were stimulated with IL-23, IL-1β, or both for 48 h, or with PMA + ionomycin for 24 h. IFN-γ levels in the supernatants were assessed by LEGENDplex multiplex ELISA. Data were normalized against levels in the absence of stimulation. Healthy (*n* = 21) controls, *IL23R*^G149R/G149R^ (*n* = 2), *IL23R*^G300V/G300V^ (*n* = 2), *IL23R*^R381Q/R381Q^ (*n* = 11), *IL12R*β*1*^−/−^ (*n* = 1), and *IL23R*^−/−^ (*n* = 3) individuals were investigated. Nonparametric Mann–Whitney *U* tests were used for analysis, with *P < 0.05, **P < 0.01, and ***P < 0.001. **(E and F)** The production of IFN-γ and TNF after PMA-ionomycin stimulation was monitored as a control, as shown in Fig. 4, D and E.

## References

1. Acosta-Rodriguez, E.V., G. Napolitani, A. Lanzavecchia, and F. Sallusto. 2007. Interleukins 1beta and 6 but not transforming growth factor-beta are essential for the differentiation of interleukin 17-producing human T helper cells. Nat. Immunol. 8:942–949. 10.1038/ni1496

2. Alodayani, A.N., A.M. Al-Otaibi, C. Deswarte, H.H. Frayha, M. Bouaziz, M. AlHelale, T. Le Voyer, A. Nieto-Patlan, V. Rattina, M. AlZahrani, et al. 2018. Mendelian susceptibility to mycobacterial disease caused by a novel founder IL12B mutation in Saudi Arabia. J. Clin. Immunol. 38:278–282. 10.1007/s10875-018-0490-2

3. Altare, F., A. Durandy, D. Lammas, J.F. Emile, S. Lamhamedi, F. Le Deist, P. Drysdale, E. Jouanguy, R. Döffinger, F. Bernaudin, et al. 1998a. Impairment of mycobacterial immunity in human interleukin-12 receptor deficiency. Science. 280:1432–1435. 10.1126/science.280.5368.1432

4. Altare, F., D. Lammas, P. Revy, E. Jouanguy, R. Döffinger, S. Lamhamedi, P. Drysdale, D. Tollner, J. Girdlestone, P. Darbyshire, et al. 1998b. Inherited interleukin 12 deficiency in a child with bacille Calmette-Guérin and Salmonella enteritidis disseminated infection. J. Clin. Invest. 102:2035–2040. 10.1172/jci4950

5. Amezquita, R.A., A.T.L. Lun, E. Becht, V.J. Carey, L.N. Carpp, L. Geistlinger, F. Marini, K. Rue-Albrecht, D. Risso, C. Soneson, et al. 2020. Orchestrating single-cell analysis with Bioconductor. Nat. Methods. 17:137–145. 10.1038/s41592-019-0654-x

6. Belkadi, A., V. Pedergnana, A. Cobat, Y. Itan, Q.B. Vincent, A. Abhyankar, L. Sha ng, J. El Baghdadi, A. Bousfiha, et al; Exome/Array Consortium. 2016. Whole-exome sequencing to analyze population structure, parental inbreeding, and familial linkage. Proc. Natl. Acad. Sci. USA. 113:6713–6718. 10.1073/pnas.1606460113

7. Bohlen, J., Q. Zhou, Q. Philippot, M. Ogishi, D. Rinchai, T. Nieminen, S. Seyedpou r, N. Parvaneh, N. Rezaei, N. Yazdanpanah, et al. 2023. Human MCTS1-dependent translation of JAK2 is essential for IFN-γ immunity to mycobacteria. Cell. 186:5114–5134.e27. 10.1016/j.cell.2023.09.024

8. Boisson-Dupuis, S., P. Bastard, V. Béziat, J. Bustamante, A. Cobat, E. Jouanguy, A. Puel, J. Rosain, Q. Zhang, S.Y. Zhang, et al.. 2024. The monogenic landscape of human infectious diseases. J. Allergy Clin. Immunol. 155:768–783. 10.1016/j.jaci.2024.12.1078

9. Boisson-Dupuis, S., J. El Baghdadi, N. Parvaneh, A. Bousfiha, J. Bustamante, J. Feinberg, A. Samarina, A.V. Grant, L. Janniere, N. El Hafidi, et al. 2011. IL-12Rβ1 deficiency in two of fifty children with severe tuberculosis from Iran, Morocco, and Turkey. PLoS One. 6. e18524. 10.1371/journal.pone.0018524

10. Boisson-Dupuis, S., N. Ramirez-Alejo, Z. Li, E. Patin, G. Rao, G. Kerner, C.K. Lim, D.N. Krementsov, N. Hernandez, C.S. Ma, et al. 2018. Tuberculosis and impaired IL-23-dependent IFN-γ immunity in humans homozygous for a common TYK2 missense variant. Sci. Immunol. 3. eaau8714. 10.1126/sciimmunol.aau8714

11. Bustamante, J. 2020. Mendelian susceptibility to mycobacterial disease: Recent discoveries. Hum. Genet. 139:993–1000. 10.1007/s00439-020-02120-y

12. Camard, L., T. Stephen, H. Yahia-Cherbal, V. Guillemot, S. Mella, V. Baillet, H. Lopez-Maestre, D. Capocefalo, L. Cantini, C. Leloup, et al. 2025. IL-23 tunes inflammatory functions of human mucosal-associated invariant T cells. iScience. 28:111898. 10.1016/j.isci.2025.111898

13. Casanova, J.-L. 2025. Human immunity. J. Hum. Immun. 1:e20250001. 10.70962/jhi.20250001

14. Casanova, Jean-Laurent. 2026. Toward a monogenic architecture of human infections: From 1996 to 2026. J. Hum. Immun. 2:e20260027. 10.70962/jhi.20260027

15. Casanova, J.-L., and L. Abel. 2002. Genetic dissection of immunity to mycobacteria: The human model. Annu. Rev. Immunol. 20:581–620. 10.1146/annurev.immunol.20.081501.125851

16. Casanova, J.-L., and L. Abel. 2022. From rare disorders of immunity to common determinants of infection: Following the mechanistic thread. Cell. 185:3086–3103. 10.1016/j.cell.2022.07.004

17. Casanova, J.-L., J.D. MacMicking, and C.F. Nathan. 2024. Interferon-γ and infectious diseases: Lessons and prospects. Science. 384. eadl2016. 10.1126/science.adl2016

18. Crowell, H.L., C. Soneson, P.L. Germain, D. Calini, L. Collin, C. Raposo, D. Malhotra, and M.D. Robinson. 2020. Muscat detects subpopulation-specific state transitions from multi-sample multi-condition single-cell transcriptomics data. Nat. Commun. 11:6077. 10.1038/s41467-020-19894-4

19. Cua, D.J., and C.M. Tato. 2010. Innate IL-17-producing cells: The sentinels of the immune system. Nat. Rev. Immunol. 10:479–489. 10.1038/nri2800

20. de Beaucoudrey, L., A. Samarina, J. Bustamante, A. Cobat, S. Boisson-Dupuis, J. Feinberg, S. Al-Muhsen, L. Janniere, Y. Rose, M. de Suremain, et al. 2010. Revisiting human IL-12Rβ1 deficiency: A survey of 141 patients from 30 countries. Medicine (Baltimore). 89:381–402. 10.1097/MD.0b013e3181fdd832

21. de Jong, R., F. Altare, I.A. Haagen, D.G. Elferink, T. Boer, P.J. van Breda Vriesman, P.J. Kabel, J.M. Draaisma, J.T. van Dissel, F.P. Kroon, et al. 1998. Severe mycobacterial and Salmonella infections in interleukin-12 receptor-deficient patients. Science. 280:1435–1438. 10.1126/science.280.5368.1435

22. Duerr, R.H., K.D. Taylor, S.R. Brant, J.D. Rioux, M.S. Silverberg, M.J. Daly, A.H. Steinhart, C. Abraham, M. Regueiro, A. Griffiths, et al. 2006. A genome-wide association study identifies IL23R as an inflammatory bowel disease gene. Science. 314:1461–1463. 10.1126/science.1135245

23. Fieschi, C., and J.L. Casanova. 2003. The role of interleukin-12 in human infectious diseases: Only a faint signature. Eur. J. Immunol. 33:1461–1464. 10.1002/eji.200324038

24. Fieschi, C., S. Dupuis, E. Catherinot, J. Feinberg, J. Bustamante, A. Breiman, F. Alt are, R. Baretto, F. Le Deist, S. Kayal, et al. 2003. Low penetrance, broad resistance, and favorable outcome of interleukin 12 receptor beta1 deficiency: Medical and immunological implications. J. Exp. Med. 197:527–535. 10.1084/jem.20021769

25. Genin, E., A. Tullio-Pelet, F. Begeot, S. Lyonnet, and L. Abel. 2004. Estimating the age of rare disease mutations: The example of triple-A syndrome. J. Med. Genet. 41:445–449. 10.1136/jmg.2003.017962

26. Gerbaux, M., F. Staels, M. Willemsen, J. Neumann, L. Bücken, L. Van Meerbeeck, W. Roosens, A. Liston, S. Humblet-Baron, and R. Schrijvers. 2025. Homozygosity for the common IL23R R381Q variant associates with increased susceptibility to chronic mucocutaneous candidiasis. Eur. J. Immunol. 55. e70002. 10.1002/eji.70002

27. Hao, Y., S. Hao, E. Andersen-Nissen, W.M. Mauck, 3rd, S. Zheng, A. Butler, M.J. Lee, A.J. Wilk, C. Darby, M. Zager, et al. 2021. Integrated analysis of multimodal single-cell data. Cell. 184:3573–3587.e29. 10.1016/j.cell.2021.04.048

28. Hunter, C.A. 2005. New IL-12-family members: IL-23 and IL-27, cytokines with divergent functions. Nat. Rev. Immunol. 5:521–531. 10.1038/nri1648

29. Itan, Y., L. Shang, B. Boisson, M.J. Ciancanelli, J.G. Markle, R. Martinez-Barricarte, E. Scott, I. Shah, P.D. Stenson, J. Gleeson, et al. 2016. The mutation significance cutoff: Gene-level thresholds for variant predictions. Nat. Methods. 13:109–110. 10.1038/nmeth.3739

30. Itan, Y., L. Shang, B. Boisson, E. Patin, A. Bolze, M. Moncada-Vélez, E. Scott, M. Ciancanelli, F.G. Lafaille, J.G. Markle, et al. 2015. The human gene damage index as a gene-level approach to prioritizing exome variants. Proc. Natl. Acad. Sci. USA. 112:13615–13620. 10.1073/pnas.1518646112

31. Kars, M.E., A.N. Başak, O.E. Onat, K. Bilguvar, J. Choi, Y. Itan, C. Caglar, R. Palv adeau, J.L. Casanova, D.N. Cooper, et al. 2021. The genetic structure of the Turkish population reveals high levels of variation and admixture. Proc. Natl. Acad. Sci. USA. 118. e2026076118. 10.1073/pnas.2026076118

32. Kerner, G., G. Laval, E. Patin, S. Boisson-Dupuis, L. Abel, J.L. Casanova, and L. Quintana-Murci. 2021. Human ancient DNA analyses reveal the high burden of tuberculosis in Europeans over the last 2,000 years. Am. J. Hum. Genet. 108:517–524. 10.1016/j.ajhg.2021.02.009

33. Kerner, G., A.L. Neehus, Q. Philippot, J. Bohlen, D. Rinchai, N. Kerrouche, A. Puel, S.Y. Zhang, S. Boisson-Dupuis, L. Abel, et al. 2023. Genetic adaptation to pathogens and increased risk of inflammatory disorders in post-Neolithic Europe. Cell Genom. 3:100248. 10.1016/j.xgen.2022.100248

34. Kerner, G., N. Ramirez-Alejo, Y. Seeleuthner, R. Yang, M. Ogishi, A. Cobat, E. Patin, L. Quintana-Murci, S. Boisson-Dupuis, J.L. Casanova, et al.. 2019. Homozygosity for TYK2 P1104A underlies tuberculosis in about 1% of patients in a cohort of European ancestry. Proc. Natl. Acad. Sci. USA. 116:10430–10434. 10.1073/pnas.1903561116

35. Khavandegar, A., S.A. Mahdaviani, M. Zaki-Dizaji, F. Khalili-Moghaddam, S. Ansari, S. Alijani, N. Taherzadeh-Ghahfarrokhi, D. Mansouri, J.L. Casanova, J. Bustamante, et al.. 2024. Genetic, immunologic, and clinical features of 830 patients with mendelian susceptibility to mycobacterial diseases (MSMD): A systematic review. J. Allergy Clin. Immunol. 153:1432–1444. 10.1016/j.jaci.2024.01.021

36. Korotkevich, G., V. Sukhov, N. Budin, B. Shpak, M.N. Artyomov, and A. Sergushichev. 2021. Fast gene set enrichment analysis. bioRxiv. 10.1101/060012 (Preprint posted February 01, 2021).

37. Lay, C.S., A. Bridges, J. Goulding, S.J. Briddon, Z. Soloviev, P.D. Craggs, and S.J. Hill. 2022. Probing the binding of interleukin-23 to individual receptor components and the IL-23 heteromeric receptor complex in living cells using NanoBRET. Cell Chem Biol. 29:19–29.e6. 10.1016/j.chembiol.2021.05.002

38. Lay, C.S., A. Isidro-Llobet, L.E. Kilpatrick, P.D. Craggs, and S.J. Hill. 2023. Characterisation of IL-23 receptor antagonists and disease relevant mutants using fluorescent probes. Nat. Commun. 14:2882. 10.1038/s41467-023-38541-2

39. Le Voyer, T., A.L. Neehus, R. Yang, M. Ogishi, J. Rosain, F. Alroqi, M. Alshalan, S. Blumental, F. Al Ali, T. Khan, et al. 2021. Inherited deficiency of stress granule ZNFX1 in patients with monocytosis and mycobacterial disease. Proc. Natl. Acad. Sci. USA. 118. e2102804118. 10.1073/pnas.2102804118

40. Liberzon, A., A. Subramanian, R. Pinchback, H. Thorvaldsdottir, P. Tamayo, and J.P. Mesirov. 2011. Molecular signatures database (MSigDB) 3.0. Bioinformatics. 27:1739–1740. 10.1093/bioinformatics/btr260

41. Love, M.I., W. Huber, and S. Anders. 2014. Moderated estimation of fold change and dispersion for RNA-seq data with DESeq2. Genome Biol. 15:550. 10.1186/s13059-014-0550-8

42. Martin-Fernandez, M., S. Buta, T. Le Voyer, Z. Li, L.T. Dynesen, F. Vuillier, L. Franklin, F. Ailal, A. Muglia Amancio, L. Malle, et al. 2022. A partial form of inherited human USP18 deficiency underlies infection and inflammation. J. Exp. Med. 219. e20211273. 10.1084/jem.20211273

43. Martinez-Barricarte, R., J.G. Markle, C.S. Ma, E.K. Deenick, N. Ramirez-Alejo, F. Mele, D. Latorre, S.A. Mahdaviani, C. Aytekin, D. Mansouri, et al. 2018. Human IFN-γ immunity to mycobacteria is governed by both IL-12 and IL-23. Sci. Immunol. 3. eaau6759. 10.1126/sciimmunol.aau6759

44. Mbatchou, J., L. Barnard, J. Backman, A. Marcketta, J.A. Kosmicki, A. Ziyatdinov, C. Benner, C. O’Dushlaine, M. Barber, B. Boutkov, et al. 2021. Computationally efficient whole-genome regression for quantitative and binary traits. Nat. Genet. 53:1097–1103. 10.1038/s41588-021-00870-7

45. Melo, K.M., F.S. Tavares, T.S. Antunes, A. Condino-Neto, G.R. Silva Segundo, A.C.T. Macedo, A.P. Ferreira, and C.F.C. Valente. 2024. Autosomal recessive IL-12p40 deficiency due to a mutation in the *IL12B* gene: Report of a Brazilian patient with lymph node mycobacterial infection. Pediatr. Allergy Immunol. Pulmonol. 37:33–36. 10.1089/ped.2022.0206

46. Momenilandi, M., R. Levy, S. Sobrino, J. Li, C. Lagresle-Peyrou, H. Esmaeilzadeh, A. Fayand, C. Le Floc’h, A. Guerin, E. Della Mina, et al. 2024. FLT3L governs the development of partially overlapping hematopoietic lineages in humans and mice. Cell. 187:2817–2837.e31. 10.1016/j.cell.2024.04.009

47. Momozawa, Y., M. Mni, K. Nakamura, W. Coppieters, S. Almer, L. Amininejad, I. Cleynen, J.F. Colombel, P. de Rijk, O. Dewit, et al. 2011. Resequencing of positional candidates identifies low frequency IL23R coding variants protecting against inflammatory bowel disease. Nat. Genet. 43:43–47. 10.1038/ng.733

48. Neehus, A.-L., B. Carey, M. Landekic, P. Panikulam, G. Deutsch, M. Ogishi, C.A. Arango-Franco, Q. Philippot, M. Modaresi, I. Mohammadzadeh, et al. 2024. Human inherited CCR2 deficiency underlies progressive polycystic lung disease. Cell. 187:390–408.e23. 10.1016/j.cell.2023.11.036

49. Ogishi, M., A.A. Arias, R. Yang, J.E. Han, P. Zhang, D. Rinchai, J. Halpern, J. Mulwa, N. Keating, M. Chrabieh, et al. 2022. Impaired IL-23-dependent induction of IFN-γ underlies mycobacterial disease in patients with inherited TYK2 deficiency. J. Exp. Med. 219. e20220094. 10.1084/jem.20220094

50. Ogishi, M., J. Puchan, R. Yang, A.A. Arias, J.E. Han, T. Nguyen, R. Gutierrez-Cozar, C. Conil, Y. Seeleuthner, D. Rinchai, et al. 2025. Human LY9 governs CD4^+^ T cell IFN-γ immunity to *Mycobacterium tuberculosis*. Sci. Immunol. 10. eads7377. 10.1126/sciimmunol.ads7377

51. Ogishi, M., R. Yang, R. Rodriguez, D.P. Golec, E. Martin, Q. Philippot, J. Bohlen, S.J. Pelham, A.A. Arias, T. Khan, et al. 2023. Inherited human ITK deficiency impairs IFN-γ immunity and underlies tuberculosis. J. Exp. Med. 220. e20220484. 10.1084/jem.20220484

52. Oppmann, B., R. Lesley, B. Blom, J.C. Timans, Y. Xu, B. Hunte, F. Vega, N. Yu, J. Wang, K. Singh, et al. 2000. Novel p19 protein engages IL-12p40 to form a cytokine, IL-23, with biological activities similar as well as distinct from IL-12. Immunity. 13:715–725. 10.1016/s1074-7613(00)00070-4

53. Parham, C., M. Chirica, J. Timans, E. Vaisberg, M. Travis, J. Cheung, S. Pflanz, R. Zhang, K.P. Singh, F. Vega, et al. 2002. A receptor for the heterodimeric cytokine IL-23 is composed of IL-12Rbeta1 and a novel cytokine receptor subunit, IL-23R. J. Immunol. 168:5699–5708. 10.4049/jimmunol.168.11.5699

54. Philippot, Q., M. Ogishi, J. Bohlen, J. Puchan, A.A. Arias, T. Nguyen, M. Martin-Fernandez, C. Conil, D. Rinchai, M. Momenilandi, et al. 2023. Human IL-23 is essential for IFN-γ-dependent immunity to mycobacteria. Sci. Immunol. 8. eabq5204. 10.1126/sciimmunol.abq5204

55. Prando, C., A. Samarina, J. Bustamante, S. Boisson-Dupuis, A. Cobat, C. Picard, Z. AlSum, S. Al-Jumaah, S. Al-Hajjar, H. Frayha, et al. 2013. Inherited IL-12p40 deficiency: Genetic, immunologic, and clinical features of 49 patients from 30 kindreds. Medicine (Baltimore). 92:109–122. 10.1097/MD.0b013e31828a01f9

56. Purcell, S., B. Neale, K. Todd-Brown, L. Thomas, M.A. Ferreira, D. Bender, J. Maller, P. Sklar, P.I. de Bakker, M.J. Daly, et al.. 2007. PLINK: A tool set for whole-genome association and population-based linkage analyses. Am. J. Hum. Genet. 81:559–575. 10.1086/519795

57. Rapaport, F., B. Boisson, A. Gregor, V. Béziat, S. Boisson-Dupuis, J. Bustamante, E. Jouanguy, A. Puel, J. Rosain, Q. Zhang, et al. 2020. Negative selection on human genes underlying inborn errors depends on disease outcome and both the mode and mechanism of inheritance. Proc. Natl. Acad. Sci. USA. 118. e2001248118. 10.1073/pnas.2001248118

58. Ritchie, M.E., B. Phipson, D. Wu, Y. Hu, C.W. Law, W. Shi, and G.K. Smyth. 2015. Limma powers differential expression analyses for RNA-sequencing and microarray studies. Nucleic Acids Res. 43:e47. 10.1093/nar/gkv007

59. Rosain, J., X.F. Kong, R. Martinez-Barricarte, C. Oleaga-Quintas, N. Ramirez-Alejo, J. Markle, S. Okada, S. Boisson-Dupuis, J.L. Casanova, and J. Bustamante. 2018. Mendelian susceptibility to mycobacterial disease: 2014-2018 update. Immunol. Cell Biol. 97:360–367. 10.1111/imcb.12210

60. Rosain, J., A.L. Neehus, J. Manry, R. Yang, J. Le Pen, W. Daher, Z. Liu, Y.H. Chan, N. Tahuil, O. Turel, et al. 2023. Human IRF1 governs macrophagic IFN-γ immunity to mycobacteria. Cell. 186:621–645.e33. 10.1016/j.cell.2022.12.038

61. Staels, F., F. Lorenzetti, K. De Keukeleere, M. Willemsen, M. Gerbaux, J. Neumann, T. Tousseyn, E. Pasciuto, P. De Munter, X. Bossuyt, et al. 2022. A novel homozygous stop mutation in IL23R causes mendelian susceptibility to mycobacterial disease. J. Clin. Immunol. 42:1638–1652. 10.1007/s10875-022-01320-7

62. Teng, M.W., E.P. Bowman, J.J. McElwee, M.J. Smyth, J.L. Casanova, A.M. Cooper, and D.J. Cua. 2015. IL-12 and IL-23 cytokines: From discovery to targeted therapies for immune-mediated inflammatory diseases. Nat Med. 21:719–729. 10.1038/nm.3895

63. Wang, R.J., S.I. Al-Saffar, J. Rogers, and M.W. Hahn. 2023. Human generation times across the past 250,000 years. Sci Adv. 9. eabm7047. 10.1126/sciadv.abm7047

64. Wilson, N.J., K. Boniface, J.R. Chan, B.S. McKenzie, W.M. Blumenschein, J.D. Mattson, B. Basham, K. Smith, T. Chen, F. Morel, et al. 2007. Development, cytokine profile and function of human interleukin 17-producing helper T cells. Nat. Immunol. 8:950–957. 10.1038/ni1497

65. Wu, M.C., S. Lee, T. Cai, Y. Li, M. Boehnke, and X. Lin. 2011. Rare-variant association testing for sequencing data with the sequence kernel association test. Am. J. Hum. Genet. 89:82–93. 10.1016/j.ajhg.2011.05.029

66. Yang, R., F. Mele, L. Worley, D. Langlais, J. Rosain, I. Benhsaien, H. Elarabi, C.A. Croft, J.M. Doisne, P. Zhang, et al. 2020. Human T-bet governs innate and innate-like adaptive IFN-γ immunity against mycobacteria. Cell. 183:1826–1847.e31. 10.1016/j.cell.2020.10.046

67. Zhu, A., J.G. Ibrahim, and M.I. Love. 2019. Heavy-tailed prior distributions for sequence count data: Removing the noise and preserving large differences. Bioinformatics. 35:2084–2092. 10.1093/bioinformatics/bty895

