## Supplementary figures and images for "Humans homozygous for rare or common hypomorphic *IL23R* variants are prone to tuberculosis"

### Supplemental Figure 1

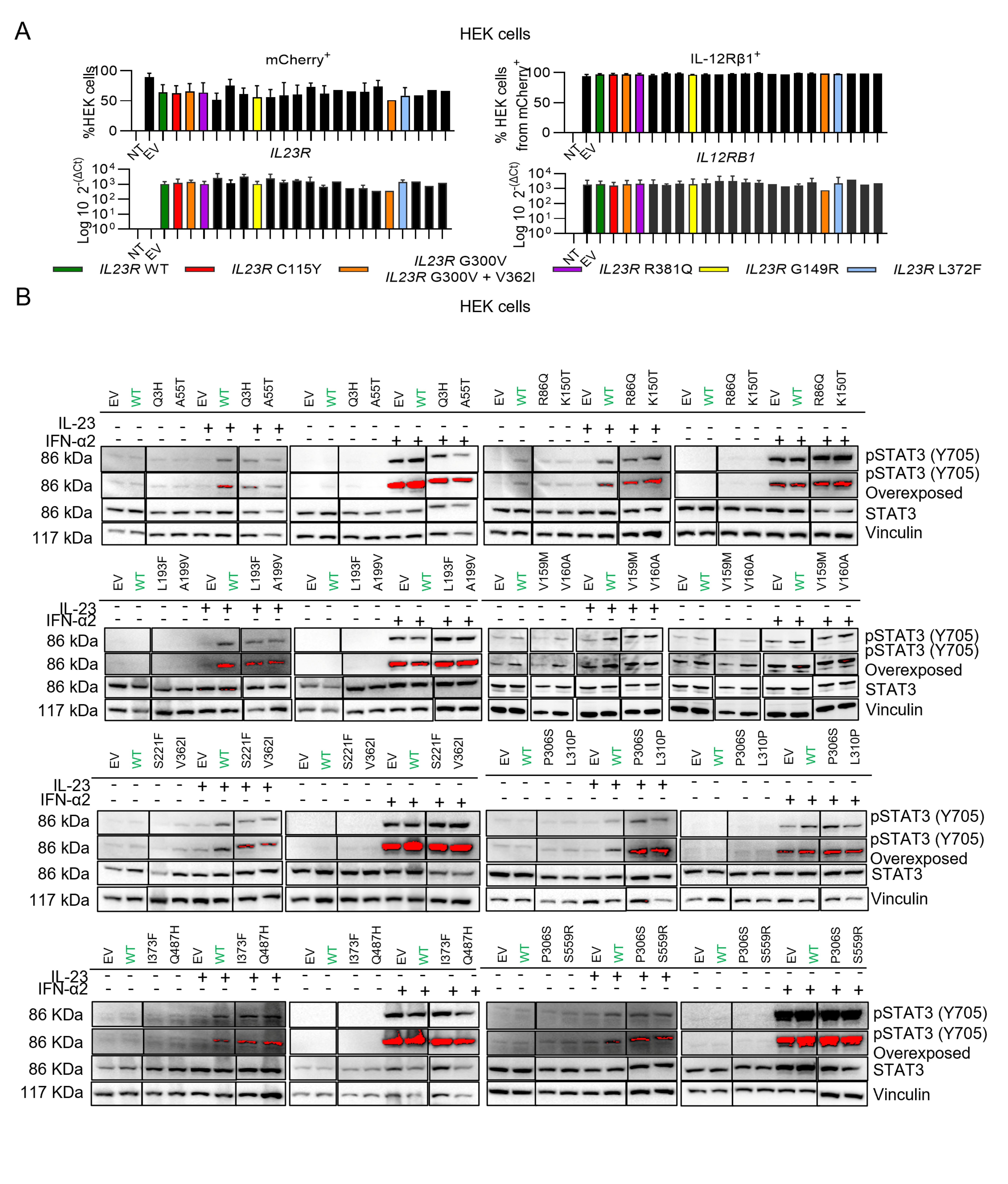

### Supplemental Figure 2

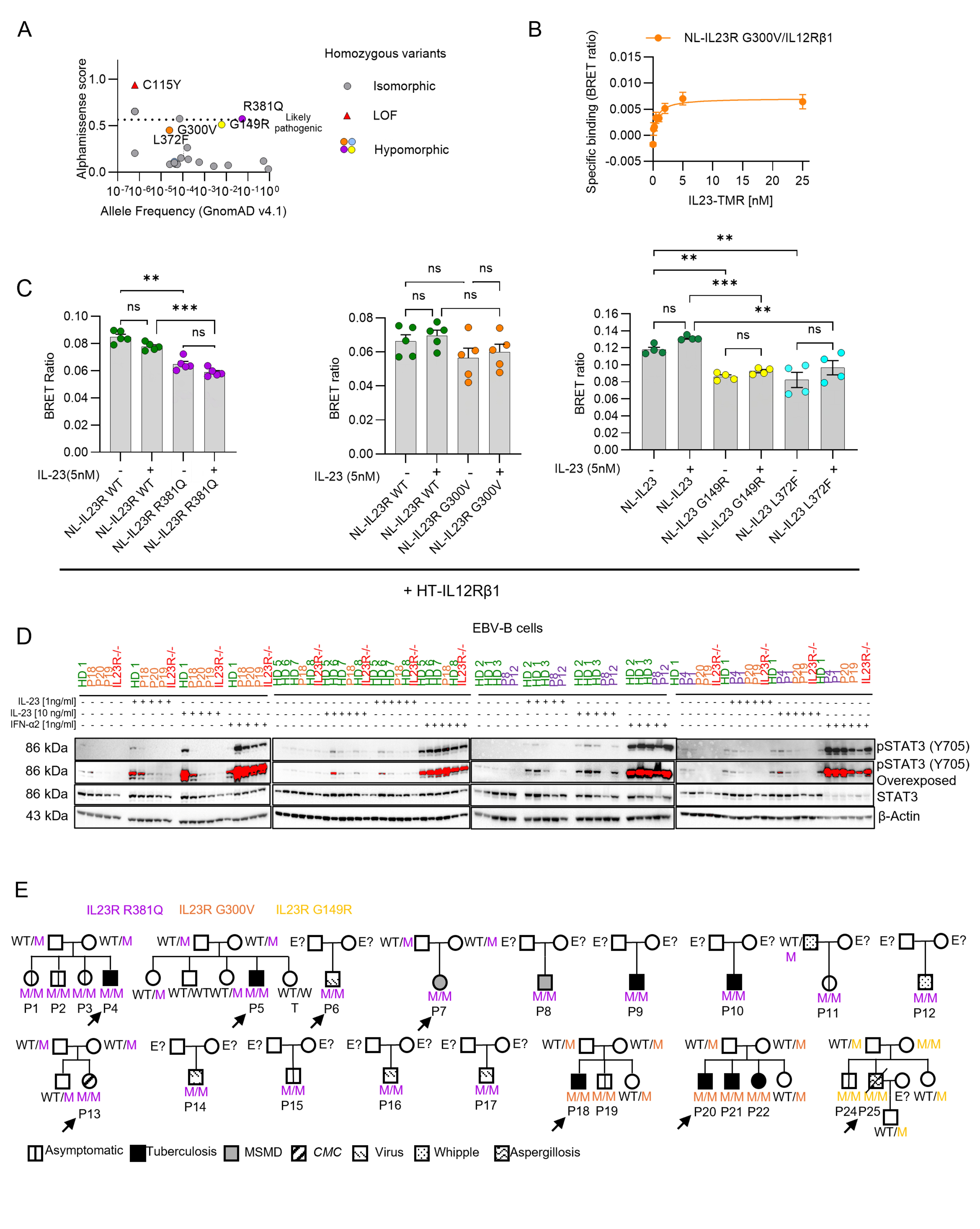

### Supplemental Figure 3

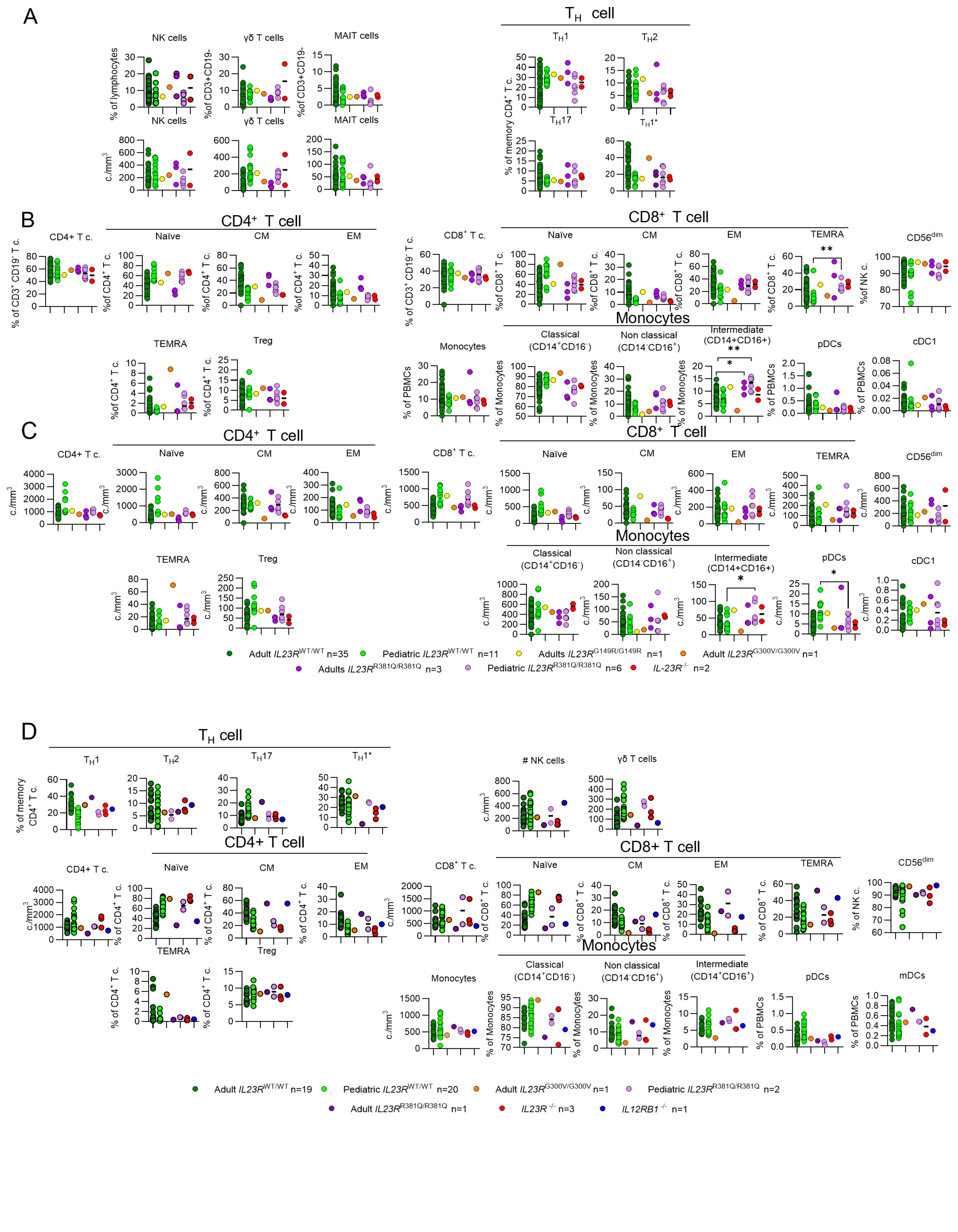

### Supplemental Figure 4

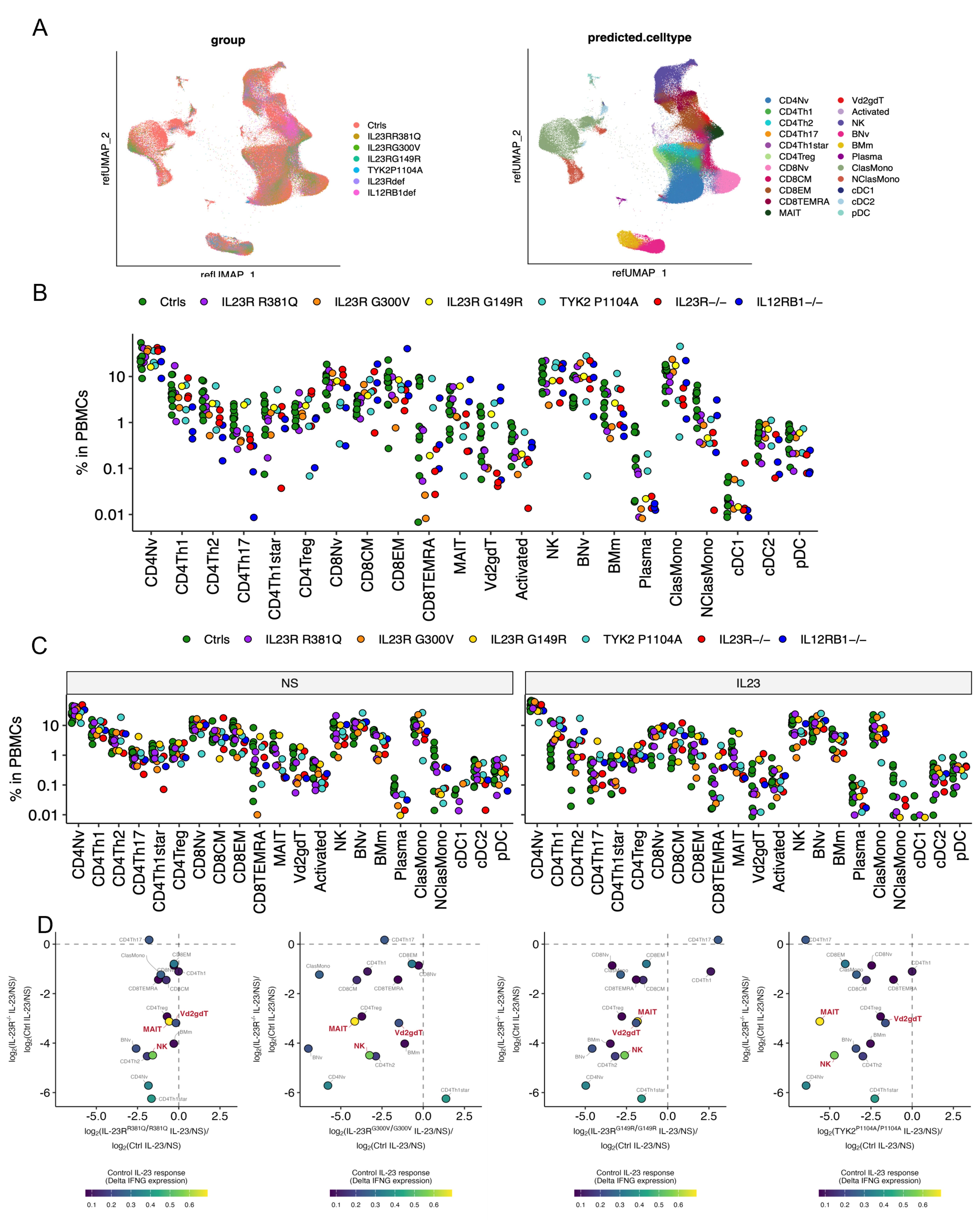

### Supplemental Figure 5

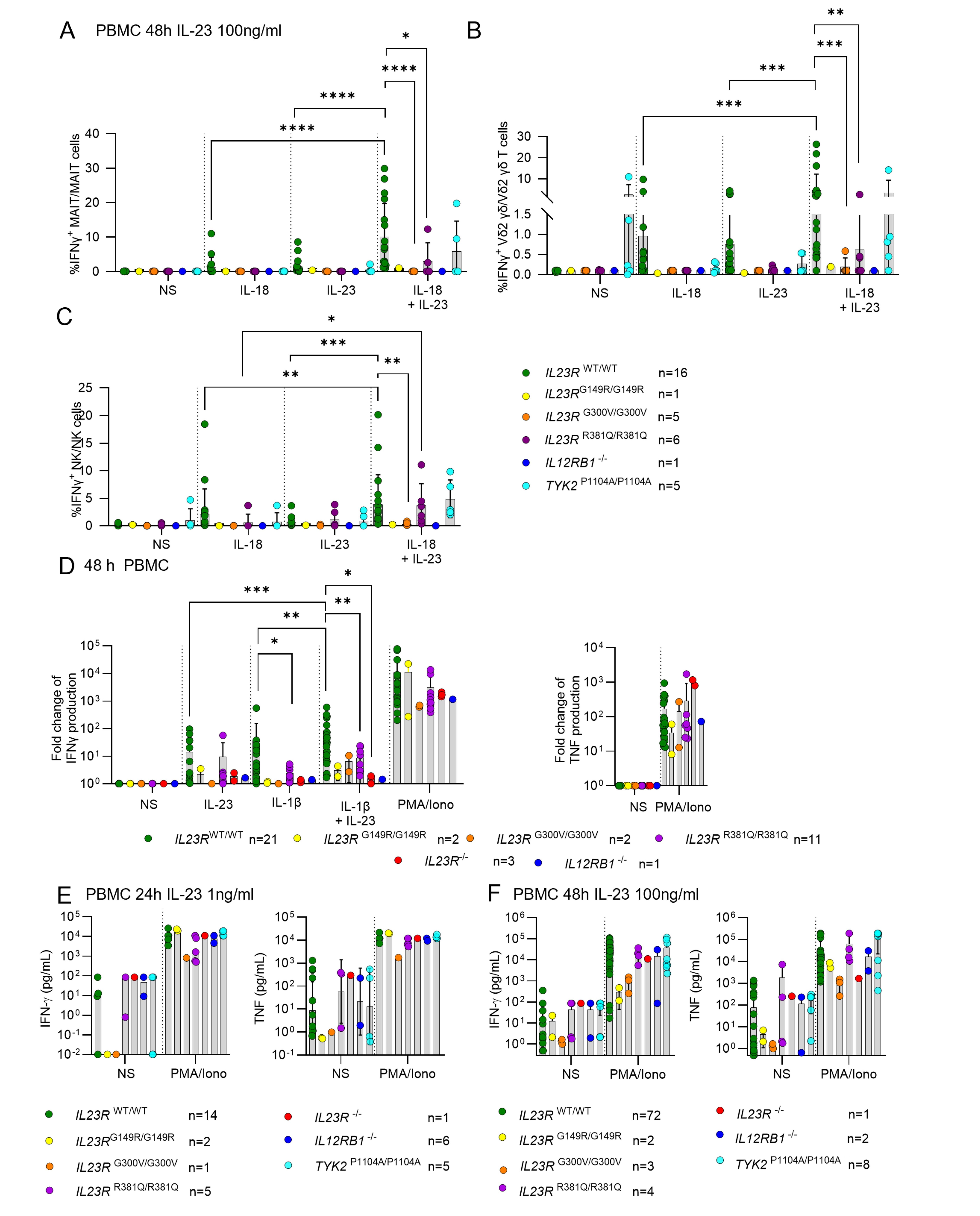
